# Metabolic engineering of *Saccharomyces cerevisiae* for second-generation ethanol production from xylo-oligosaccharides and acetate

**DOI:** 10.1101/2023.02.04.527128

**Authors:** Dielle Pierotti Procópio, Jae Won Lee, Jonghyeok Shin, Robson Tramontina, Patrícia Felix Ávila, Lívia Beatriz Brenelli, Fabio Márcio Squina, André Damasio, Sarita Cândida Rabelo, Rosana Goldbeck, Telma Teixeira Franco, David Leak, Yong-Su Jin, Thiago Olitta Basso

## Abstract

Simultaneous intracellular depolymerization of xylo-oligosaccharides (XOS) and acetate fermentation by engineered *Saccharomyces cerevisiae* offers an advance towards more cost-effective second-generation (2G) ethanol production. As xylan is one of the most abundant polysaccharides present in lignocellulosic residues, the transport and breakdown of XOS in an intracellular environment might bring a competitive advantage for recombinant strains in competition with contaminating microbes, which are always present in fermentation tanks; furthermore, acetic acid is a ubiquitous toxic component in lignocellulosic hydrolysates, deriving from hemicellulose and lignin breakdown. In the present work, the previously engineered *S. cerevisiae* strain, SR8A6S3, expressing NADPH-linked xylose reductase (XR), NAD^+^-linked xylitol dehydrogenase (XDH) (for xylose assimilation), as well as NADH-linked acetylating acetaldehyde dehydrogenase (AADH) and acetyl-CoA synthetase (ACS) (for an NADH-dependent acetate reduction pathway), was used as the host for expressing of two β-xylosidases, *GH43-2* and *GH43-7*, and a xylodextrin transporter, *CDT-2*, from *Neurospora crassa*, yielding the engineered strain SR8A6S3-CDT_2_-GH43_2/7_. Both β-xylosidases and the transporter were introduced by replacing two endogenous genes, *GRE3* and *SOR1*, that encode aldose reductase and sorbitol (xylitol) dehydrogenase, respectively, which catalyse steps in xylitol production. Xylitol accumulation during xylose fermentation is a problem for 2G ethanol production since it reduces final ethanol yield. The engineered strain, SR8A6S3-CDT_2_-GH43_2/7_, produced ethanol through simultaneous co-utilization of XOS, xylose, and acetate. The mutant strain produced 60% more ethanol and 12% less xylitol than the control strain when a hemicellulosic hydrolysate was used as a mono- and oligosaccharide source. Similarly, the ethanol yield was 84% higher for the engineered strain using hydrolysed xylan compared with the parental strain. The consumption of XOS, xylose, and acetate expands the capabilities of *S. cerevisiae* for utilization of all of the carbohydrate in lignocellulose, potentially increasing the efficiency of 2G biofuel production.

**Highlights:**

- Integration of XOS pathway in an acetate-xylose-consuming *S. cerevisiae* strain;
- Intracellular fermentation of XOS, acetate and xylose improved ethanol production;
- Deletion of both *sor1*Δ and *gre3*Δ reduced xylitol production.

## 1. INTRODUCTION

The production of fuel ethanol from sugarcane is a major contributor to the ongoing transition from fossil to renewable fuels and chemicals (Karp et al., 2021). To increase production without a massive increase in land use requires intensification by utilising all of the fermentable carbohydrate in the sugarcane, including that present in bagasse and straw, which is composed of lignocellulose (Raj et al., 2022; Raud et al., 2019). Successfully accessing and fermenting this fraction would also open up routes to using other agricultural by-products, such as straw and forestry residues. Their procurement cost is relatively low, besides being an abundant non-food feedstock (Ko and Lee, 2018). The lignocellulosic biofuel production process requires the deconstruction of biomass into fermentable sugars and the conversion of sugars to biofuels (Li et al., 2019). Due to the complex integration of cellulose, hemicellulose, and lignin in the structure of lignocellulose, harsh pre-treatment is required to access the carbohydrate polymers for enzymatic hydrolysis and fermentation, which can result in the production of by-products such as furans, organic acids, phenols and inorganic salts which can inhibit microbial metabolism (Ask et al., 2013; Kłosowski and Mikulski, 2021; Tramontina et al., 2020).

Pre-treatment aims to reduce the crystallinity of cellulose, and partially degrade hemicellulose and lignin to increase the susceptibility of the biomass to enzymatic cocktails, which in turn are necessary to breakdown polysaccharides into fermentable monomeric sugars (Kłosowski and Mikulski, 2021; Sarkar et al., 2012; Sharma et al., 2020). However, during the degradation of hemicellulose and lignin, acetic acid production is unavoidable as hemicellulose and lignin are acetylated (Chen et al., 2019; Klinke et al., 2004). This is toxic to yeast metabolism, reducing sugar fermentation efficiency and biofuel yield (Almeida et al., 2007; Kłosowski and Mikulski, 2021; Salas-Navarrete et al., 2022). Weak organic acids, such as acetic acid, can diffuse undissociated through the cell membrane and dissociate inside the cell, releasing protons and lowering the internal pH value (Bellissimi et al., 2009; Kłosowski and Mikulski, 2021). To overcome the inhibitory effect of acetic acid, Zhang et al., (2016) introduced an optimized route for acetate reduction, through the expression of three copies of codon-optimized acetaldehyde dehydrogenase - *adhE* (*CO_adhE*) from *Escherichia coli*, and three copies of a mutated acetyl-CoA synthetase - *ACS* (*ACS*Opt*) from *Salmonellas enterica* into a xylose-fermenting *S. cerevisiae* strain, which produces recombinant NADPH-linked xylose reductase (XR) and NAD^+^-linked xylitol dehydrogenase (XDH), yielding strain SR8A6S3. This strategy enabled efficient xylose fermentation with 29.7% higher ethanol yield and 70.7% lower by-product (xylitol and glycerol) production when cultivated in YP medium supplemented with 20 g L^-1^ glucose, 80 g L^-1^ xylose, and 8 g L^-1^ acetate under strict anaerobic (anoxic) conditions. The reduction of acetate to ethanol serves as an electron sink to alleviate the redox cofactor imbalance resulting from *XR* and *XDH* activities (Wei et al., 2013), with NAD^+^ generated from the reductive metabolism of acetate being available for XDH activity, thus reducing the production of xylitol and glycerol. Thus, this strategy can provide multiple benefits for the ethanol industry (Zhang et al., 2016).

Although SR8A6S3 can tolerate acetic acid present in lignocellulosic hydrolysates, many other inhibitory compounds are also released during the pre-treatment steps (Kłosowski and Mikulski, 2021). While less severe pre-treatment could be considered for achieving a lower concentration of inhibitors, a large amount of cellulase and hemicellulase enzyme cocktails would still be required for converting cellulose and hemicellulose into monomeric sugars, posing unsolved economic and logistical challenges for the industry (Adsul et al., 2020; Chundawat et al., 2011; Himmel et al., 2007; Li et al., 2015). One possible strategy to achieve economic 2G ethanol is to use *S. cerevisiae* strains genetically modified to transport and intracellularly utilize cellulose and hemicellulose-derived oligosaccharides. Such a microorganism might have a competitive advantage over other microorganisms, such as contaminating bacteria and wild *Saccharomyces* and non-*Saccharomyces* species, which are not able to metabolize oligosaccharides, as well as requiring lower amounts of hemi/cellulolytic enzymes, which should translate into a cheaper process (Procópio et al., 2022).

In previous work, Li and co-authors (Li et al., 2015) expanded xylose utilization by an engineered *S. cerevisiae* strain, to incorporate the transport and intracellular hydrolysis of XOS to xylose monomers through the expression of two β-xylosidases, *GH43-2*, and *GH23-7*, and a XOS-transporter, *CDT-2*, from *N. crassa* in a xylose-utilizing host strain. Both glycoside hydrolases (GH) catalyse the hydrolysis of 1,4-β-D-xylosidic linkages in xylan (Mewis et al., 2016). The new strain could produce more than 30 g L^-1^ of ethanol in 72h of cultivation in an optimized minimum medium (oMM) supplemented with 4% xylose and 3% XOS under anaerobic conditions.

In this work, we used the SR8A6S3 strain as a platform for the construction of a yeast strain able to ferment XOS, xylose, and acetate into ethanol (**Fig. 1**). Genes encoding the XOS-transporter (c*dt-2*) and both of the β-xylosidases (*gh43-2* and *gh43-7*) from *N. crassa* were integrated into the SR8A6S3 genome (highlighted in the yellow area in **Fig. 1**). First, a high expression cassette for *cdt-2* expression was integrated into the sorbitol (xylitol) dehydrogenase *locus*, encoded by gene *sor1* gene, through the locus-specific CAS-9-based integration system (Stovicek et al., 2015). Then, both *gh43-2* and *gh43-7* under the control of the *GAP* and *CCW12* promoters, respectively, were integrated into the aldose reductase encoded by *gre3* gene using the same locus-specific integration tool. The resulting disruption of *GRE3* and *SOR1* was designed to mitigate xylitol production and divert more carbon towards ethanol production in the recombinant strain (Jeong et al., 2020; Toivari et al., 2004; Träff et al., 2001). Conversion of hemicellulosic-derived residues into industrial products, such as 2G ethanol, can contribute to the progress of global warming mitigation (Sun et al., 2021).

**Fig. 1.**
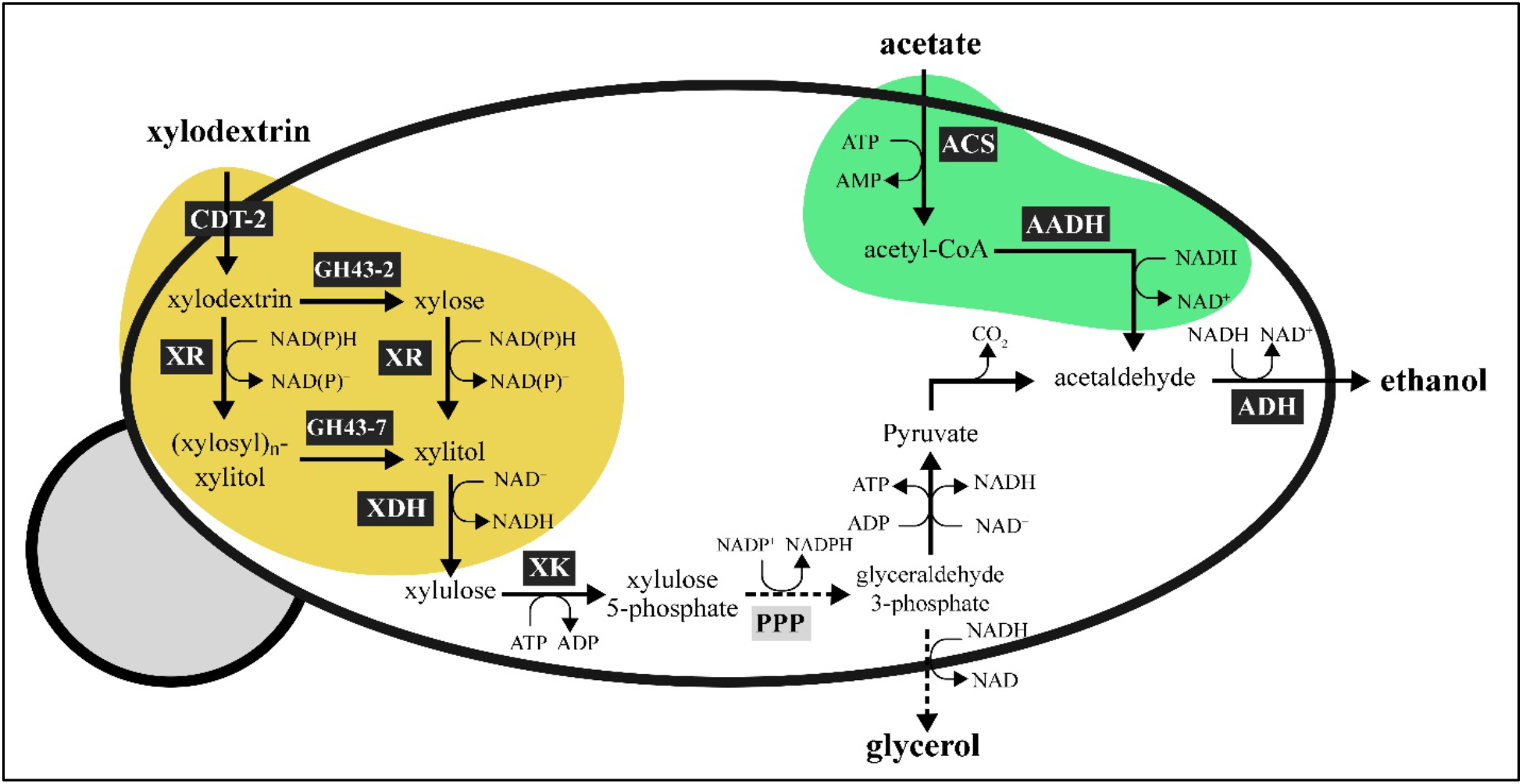
Expected routes of XOS metabolism after expression of the XOS-transporter (*CTD-2*) and beta-xylosidases (*GH43-2* and *GH43-7*) from *N. crassa* in SR8A6S3, including xylose metabolism by xylose reductase (XR) and xylitol dehydrogenase (XDH) from *S. stipitis*. The surplus NADH produced during xylose fermentation can be exploited to detoxify acetate, reducing it to ethanol through the exogenous acetate reduction pathway, involving conversion of acetate into acetyl-CoA by acetyl CoA synthetase (ACS), production of acetaldehyde from the acetyl-CoA by the acetylating acetaldehyde dehydrogenase (AADH) and ethanol production from acetaldehyde by the action of alcohol dehydrogenase (ADH).

## 2. MATERIALS AND METHODS

### 2.1. Strains and media

*E. coli* strain DH5α was used for the construction and propagation of plasmids. *E. coli* was cultured in Lysogeny Broth (LB) medium (5 g L^-1^ yeast extract, 10 g L^-1^ tryptone, and 10 g L^-1^ NaCl) at 37 °C and 100 μ/mL ampicillin (LBA) was added for selection when required. All engineered *S. cerevisiae* strains used and constructed in this work are summarized in **Table 1**. Yeast strains transformed with plasmids containing antibiotics were propagated on YPD plates supplemented with the plasmid corresponding antibiotics, such as clonNAT (100 μg mL^-1^), geneticin G418 (200 μg mL^-1^), hygromycin B (200 μg mL^-1^). The SR8A6S3-CDT_2_ strain was generated by integrating the CDT-2 transporter overexpressing gene cassette into the *SOR1* locus of the SR8A6S3 genome. To construct an XOS-utilizing strain the _p_GAP*-* GH43-7-_T_CYC-_p_CCW12-GH43-2-_T_CYC1 was integrated at the *GRE3* locus of SR8A6S3-CDT_2_, yielding strain SR8A6S3-CDT_2_-GH43_2/7_.

**Table 1.**
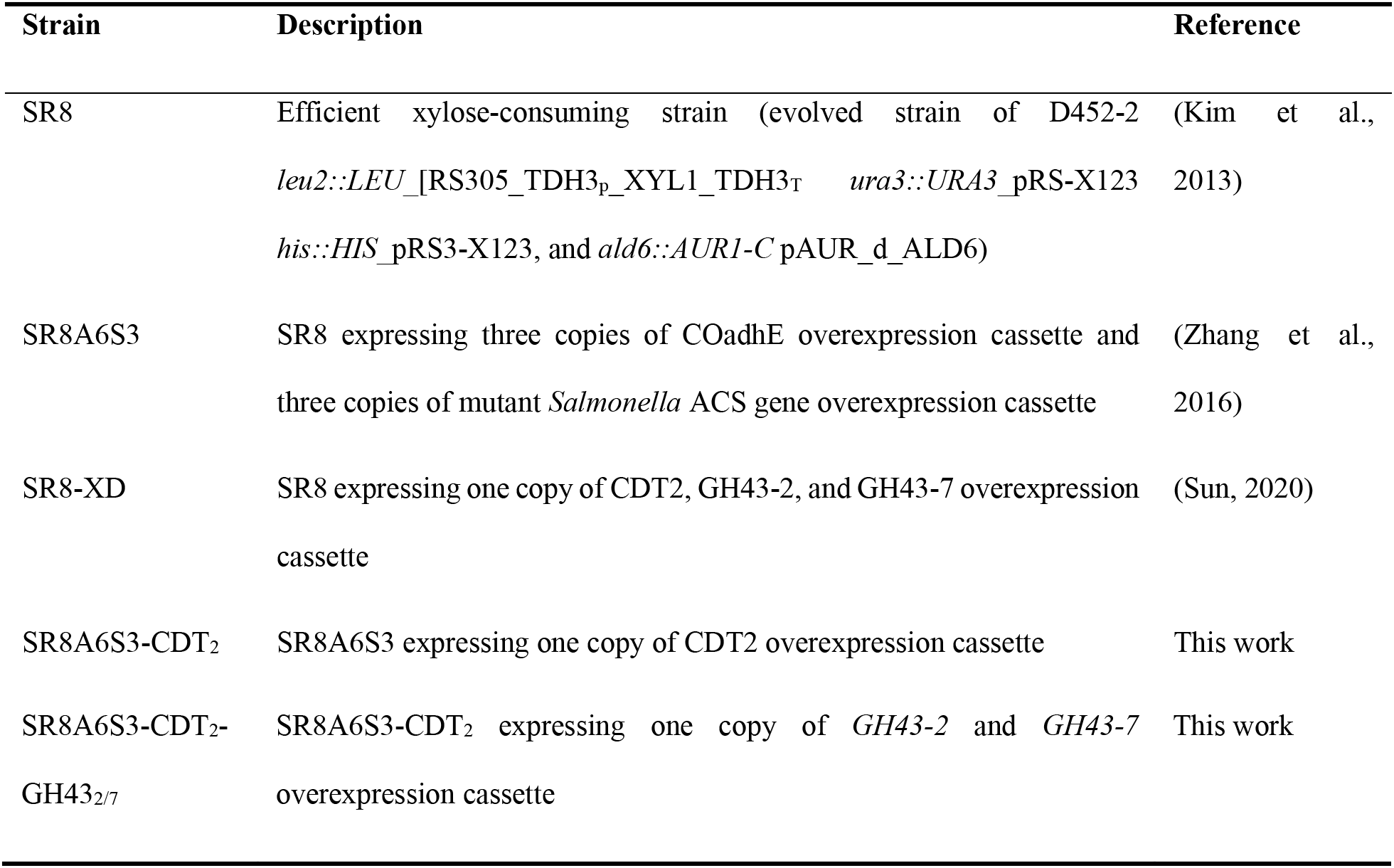
The yeast strains used in this study.

### 2.2. Plasmids and strain construction

All plasmids and primers in this work are summarized in **Tables 2** and **S1**, respectively. The guide RNA (gRNA) plasmids (**Table 3**) gRNA-sor-K and gRNA-gre-K were amplified from Cas9-NAT by using primers pair DPO_089 and DPO-090, DPO_087 and DPO_088 carrying a 20 bp PAM sequence for *SOR1* and *GRE3 loci*, respectively. The gRNAs were predicted by the website: https://www.atum.bio/eCommerce/cas9/input. All gRNA sequences are listed in **Table 3.**

**Table 2.** Plasmids used in this study.

| Plasmids | Description | Reference |
| --- | --- | --- |
| pRS42K | pRS42K, Kanamycin resistance gene | (Liu et al., 2016) |
| p426GPD | pRS426-*pTDH3-tCYC1* | Addgene (#14156) |
| Cas9-NAT | P414-*pTEF1-Cas9-tCYC1-NAT1* | Addgene (#64329) |
| p426-CDT2 | pRS425-*pPGK1-CDT2-tCYC1* | (Kim et al., 2014) |
| gRNA-sor-K | pRS42K carrying *SOR1* disruption gRNA cassette | This work |
| gRNA-gre-K | pRS42K carrying *GRE3* disruption gRNA cassette | This work |
| p426-GH43<sub>2/7</sub> | pRS426-*pTDH3-GH43-7-tCYC-pCCW12-GH43-2 tCYC1* | This work |

**Table 3.**
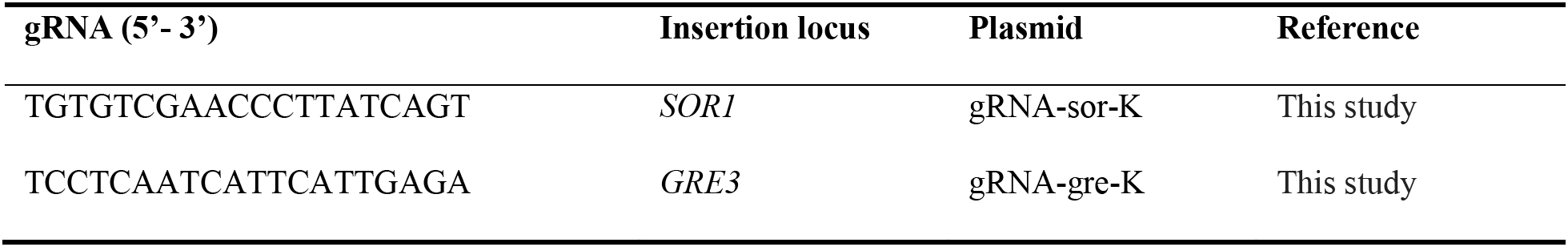
gRNA used in this study.

For genomic integration of *CDT-2* through CRISPR-Cas9-based integration in the *SOR1* gene site of SR8A6S3, CDT-2 donor DNA was amplified from plasmid pRS426-CDT_2_ using a primer pair DPO_081 and DPO_082. Transformants with *CDT-2* integration were identified by PCR using primers DPO_083 and DPO_084 and the resulting strain was designated as the SR8A6S3-CDT_2_ (**Table 1**). The PCR reaction was performed using 1.25 μL forward primer, 1.25 μL reverse primer, DNA sample 1 μL, Phusion high-fidelity DNA polymerase master mix with HF buffer (New England BioLabs) 12.5 μL, and nuclease-free water 9 μL.

To generate transformant strains expressing the GH43-7_GH32-2 gene cassette, the sequence *GH43-7-_T_CYC-_P_CCW12-GH43-2* was amplified from the genomic DNA of the XOS-consuming strain, SR8-XD (**Table S1**). Firstly, SR8-XD genomic DNA was prepared with the Rapid Yeast Genomic DNA Extraction Kit (Bio Basic Inc., Markham Ontario, CA) and quantified by NanoDrop ND-1000. Primer pairs of DPO_059 and DPO_063 were used to amplify the *GH43-7-_T_CYC-_P_CCW12-GH43-2* gene sequence. The PCR product *GH43-7-_T_CYC-_P_CCW12-GH43-2* was amplified again using a primer pair of DPO_062 and DPO_074 which has homology with plasmid p426GPD. Similarly, the plasmid p426GPD was amplified using a primer pair of DPO_064 and DPO_065. PCR was performed using 1.25 μL forward primer, 1.25 μL reverse primer, DNA sample 1 μL, Phusion high-fidelity DNA polymerase master mix with HF buffer (New England BioLabs) 12.5 μL, and nuclease-free water 9 μL. Both the linear sequences were transformed into competent *E. coli* DH5α to form the plasmid p426-GH43_2/7_ (Kostylev et al., 2015) (**Table S1**). *_P_TDH3-GH43-7-_T_CYC-_P_CCW12-GH43-2-_T_CYC1* donor DNA was amplified from plasmid p426-GH43_2/7_ using a primer pair DPO_057 and DPO_058.

Transformation of yeast cells was carried out by the polyethylene glycol (PEG)-LiAc method (Gietz et al., 1995). One microgram of DNA was used for Cas9 or gRNA plasmid transformation, 1.5 μg of donor DNA was used for homologous recombination. Correct integration was confirmed by PCR using primers DPO_069 and DPO_070. The recombinant strain was designated as SR8A6S3-CDT_2_-GH43_2/7_ (**Table 1**).

### 3.3. Enzyme activity assay and protein quantification

SR8A6S3-CDT_2_-GH342/7 and SR8A6S3 were grown in 22 mL of yeast extract-peptone (YP) medium (10 g L^-1^ yeast extract, 20 g L^-1^ peptone) containing 2% glucose, 8% xylose, and 0.8% acetate (YPDXA) until late log phase before harvesting by centrifugation. Yeast cell pellets, 0.24 g for SR8A6S3-CDT_2_-GH34_2/7_ and 0.21 g for SR8A6S3, were resuspended in buffer containing 0.1 mM CaCl_2_, 50 mM Tris-HCl, 100 mM NaCl, 1 mM DTT, 0.1% Triton X, pH 7.4, 0.1mM PMSF (Thermo Fisher Scientific). The cells were disrupted by agitation using 1 g glass beads and ultrasonic bath at 40% amplitude for 5 minutes on ice. The resulting lysates were centrifugated at 14,000×g for 20 min at 4 °C, and the clarified supernatant was used as an enzyme source for β-xylosidase assays.

β-xylosidase activity was measured according to Tramontina et al. (2016). Briefly, 30 μL of the clarified supernatant and 50 μL of 5 mM ρ-Nitrophenyl-β-D-xylopyranoside (pNPX) solution were added to 20 μL of reaction buffer (250 mM MES, and 5 mM CaCl_2_, pH 7), which was then incubated at 30 °C for 60 min for the enzyme reaction. The reaction was stopped by adding 100 μL of 2 M Na_2_CO_3_ and the amount of ρ-Nitrophenol produced was estimated spectrophotometrically at a wavelength of 405 nm and the absorbance converted to concentration using a standard curve. One unit of enzyme activity was defined as the amount of enzyme catalysing the hydrolysis of 1 μmol pNPX per minute in 1 mL of yeast intracellular lysate (μmol mL^-1^ min^-1^) “U mL^-1^”, or per mg of total lysate protein (μmol mg^-1^ min^-1^) “U mg^-1^”, or per gram of cells (μmol gcDw^-1^ min^-1^) under the described assay conditions. The protein concentrations in each sample were determined using the Bradford dye method (Bradford, 1976).

### 3.4. Fermentation and analytical methods

Anaerobic batch fermentation experiments were performed in 100 mL serum bottles with 30 mL fermentation media. Serum bottles were sealed with a butyl rubber stopper and then flushed with nitrogen gas, which had been passed through a heated, reduced copper column to remove traces of oxygen. Micro-aerobic batch fermentation experiments were performed in a 125 mL Erlenmeyer flask with 30 mL of fermentation media. Both anaerobic and micro-aerobic cultures were incubated in a rotary shaker at 100 rpm at 30 °C.

For all cultivations, yeasts were pre-grown in yeast extract-peptone (YP) medium (10 g L^-1^ yeast extract, 20 g L^-1^ peptone) supplemented with 20 g L^-1^ glucose, harvested by centrifugation at 3,134 ×g, at 4°C for 5 min, and washed three times with sterile distilled water. Washed yeast cells were inoculated in serum bottles or Erlenmeyer flasks containing either: YP supplemented with a mixture of glucose, xylose, and acetate (YPDXA); hemicellulosic hydrolysate (YPH); hemicellulosic hydrolysate, xylose and acetate (YPXAH); hydrolysed xylan (YPXy); hydrolysed xylan and acetate (YPAXy). Initial cell concentration varied according to the cultivation, OD_600_ was 1 or 10. Xylan hydrolysis was carried out according to (Ávila et al., 2020). The hemicellulosic hydrolysate from sugarcane straw was obtained by a two-stage procedure: mild acetylation at 60 °C, 30 min, 0.8% (w w^-1^) of NaOH and 10% (w w^-1^) of solids followed by hydrothermal pre-treatment at 190 °C, 20 min, 10% (w w^-1^) of solids. The hemicellulosic hydrolysate obtained after the second step was enzymatically treated with a GH11 from *Neocallimastix patriciarum* (Megazyme^®^ Ireland) as detailed described elsewhere (Brenelli et al., 2020). Afterwards, the hemicellulosic hydrolysate rich in XOS was concentrated approximately 5-fold in a rotary vacuum evaporator. **Table 4** shows the chemical composition of the XOS-rich hemicellulosic hydrolysate.

**Table 4.** Chemical composition of the XOS-rich hemicellulosic hydrolysate after treatment with a endoxylanase GH11 and concentration.

| Component | AA | FA | FT | AR | AOS | GOS | Xyl | HMF | FL | XOS |
| --- | --- | --- | --- | --- | --- | --- | --- | --- | --- | --- |
| Concentration (g L<sup>-1</sup>) | 0.77 | 2.51 | 6.40 | 3.93 | 2.45 | 6.91 | 2.74 | 0.07 | 0.01 | 49.67 |

Legend: AA - acetic acid, FA - formic acid, FT – total phenolics, AR - arabinose, AOS - arabinooligosaccharides, GOS - gluco-oligosaccharides, Xyl - xylose, HMF - hydroxymethylfurfural, FL - furfural, XOS - total xylo-oligosaccharides.

Samples were taken using syringe and needle from serum bottles or manual single-channel pipette (Gilson, USA) from Erlenmeyer flasks at appropriate intervals to measure cell growth and metabolites concentrations. Cell growth was monitored as the optical density at 600 nm (OD600) measured using a UV-visible Spectrophotometer (Biomate 5). The samples were centrifuged at 14,000×g for 10 min and supernatants diluted appropriately for the determination of glucose, xylose, xylitol, glycerol, succinate, acetic acid, and ethanol by high-performance liquid chromatography (HPLC, Agilent Technologies 1200 Series) equipped with a refractive index detector (RID). Chromatography was done on a Rezex ROA-Organic Acid H+ (8%) column (Phenomenex Inc., Torrance, CA) maintained at 60 °C, with 0.005 N H_2_SO_4_ as eluent at a flow rate of 0.6 mL min^-1^. Analyte concentrations were determined by using the RID detector.

#### 2.3.1. Xylo-oligosaccharide quantification

The enzymatic products were analysed by high-performance anion-exchange chromatography with pulsed amperometry detection (HPAEC–PAD) to detect xylose and XOS produced by the xylanase enzymes. Separation was performed using a Dionex ICS-3000 instrument (Thermo Fisher Scientific, Sunnyvale, CA, USA) with a CarboPac PA100 column (4×250 mm) and CarboPac PA100 guard column (4×50 mm), eluted with a linear gradient of A (NaOH 500 mM) and B (NaOAc 500 mM, NaOH 80 mM). The gradient program was 15 % of A and 2 % of B for 0–10 min, followed by 15–50 % of A and 2–20 % of B from 10–20 min, with a flow rate of 1.0 mL min^−1^. The integrated peak areas were converted to concentrations based on standards (×1 to×6).

## 3. RESULTS AND DISCUSSION

### 3.1. Cas9-based integration of CDT-2 expression cassette into the *SOR1* locus

Although xylitol has a variety of uses in the food, cosmetic, nutraceutical, and pharmaceutical industries (Queiroz et al., 2022), this metabolite may face competition from an available carbon source, reducing the efficiency of ethanol production. *S. cerevisiae* strains possess genes encoding enzymes capable of xylose reduction, such as *GRE3, GCY1, YPR1, YDL124W, YJR096W*, and xylitol oxidation such as *XYL2, SOR1, SOR2, XDH1*, which can result in xylitol formation during xylose fermentation (Wenger et al., 2010). To reduce xylitol production and divert the carbon to ethanol production, *SOR1* was replaced by a *CDT-2* expression cassette in the genomic DNA of strain SR8A6S3, yielding SR8A6S3-CDT_2_. The required integration of the *CDT-2* cassette was confirmed by PCR analysis. Colony PCR was performed directly from 27 colonies of the positive-control plate (**Fig. S1**). Once the desired integration was confirmed, both SR8A6S3-CDT_2_ and SR8A6S3 strains were compared in anaerobic and micro-aerobic batch cultures (**Fig. 2.A**, **2.B** and **3**) in YPDXA containing 20 g L^-1^ glucose, 80 g L^-1^ xylose, and 8 g L^-1^ acetate, with an initial OD600 of 1.

**Fig. 2.**
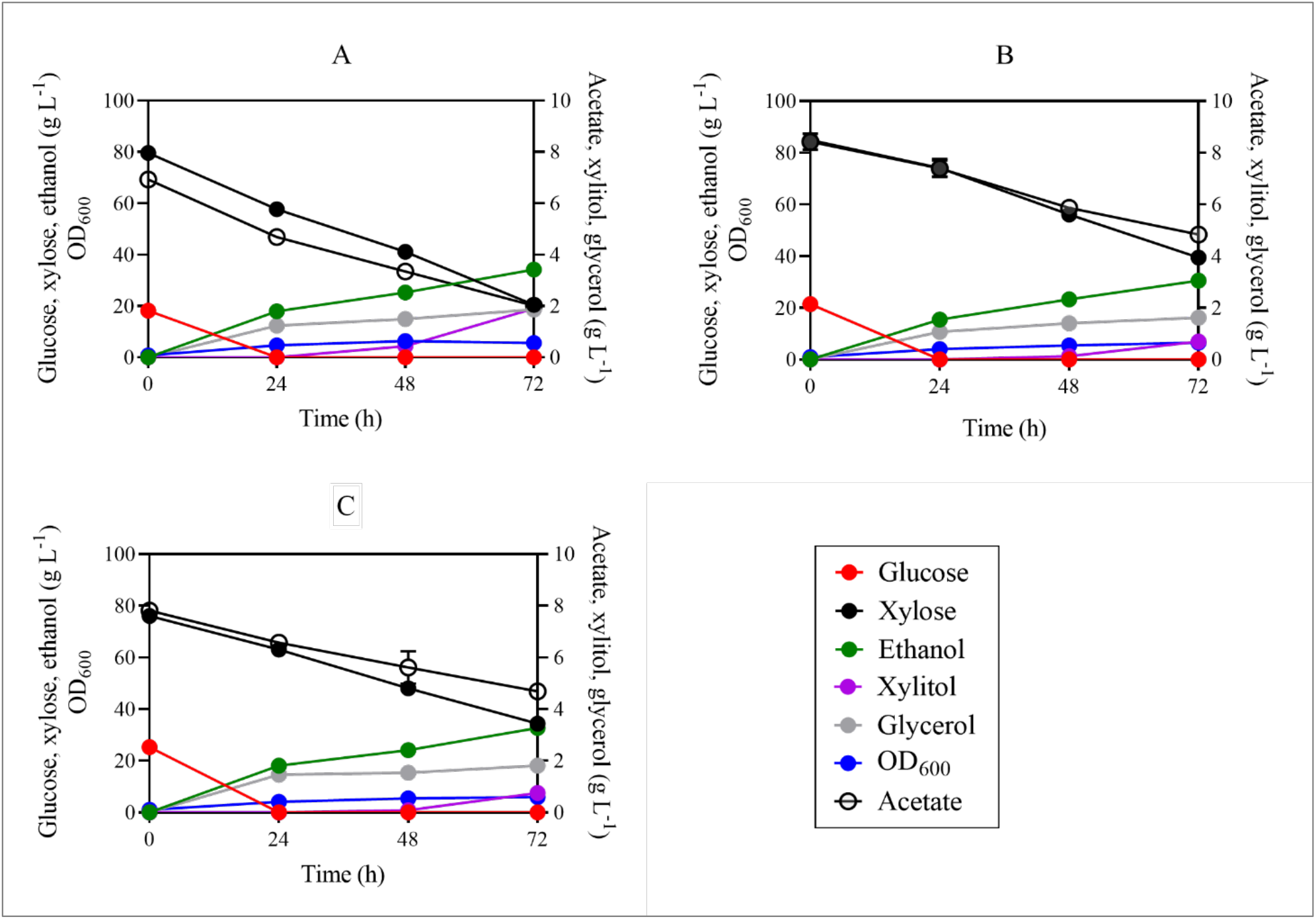
Fermentation profiles of SR8A6S3 (A), SR8A6S3-CDT_2_ (B), and SR8A6S3-CDT_2_-GH43_2/7_ (C) when fermenting YP supplemented with 20 g L^-1^ glucose, 80 g L^-1^ xylose, and 8 g L^-1^ acetate) under strictly anaerobic conditions. Data are presented as mean values and standard deviations of three independent biological replicates.

Deletion of *sor1* led to a reduced rate of xylose and acetate consumption under both anaerobic and micro-aerobic conditions (**Fig. 2.A**, **2.B,** and **3**). Under anaerobic batch cultivation, 75% of the initial xylose was consumed by the SR8A6S3 strain, while SR8A6S3-CDT_2_ was only able to consume 53% of the original concentration within 72 h (**Fig. S2.A**). The xylose consumption rate of SR8A6S3 was also higher after 24h of anaerobic cultivation in comparison to SR8A6S3-CDT_2_ (**Table 5**).

**Fig. 3.**
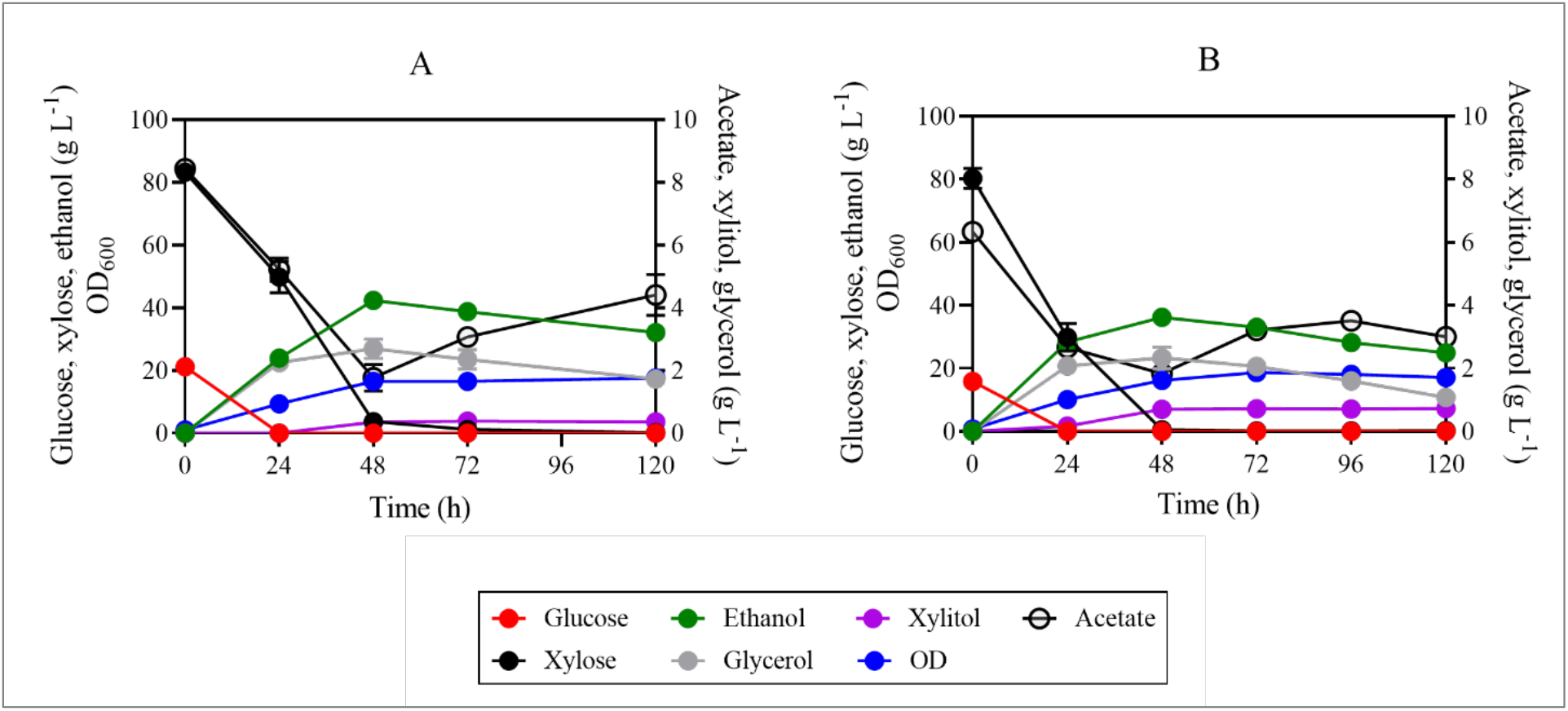
Fermentation profiles of the SR8A6S3-CDT_2_ (A), SR8A6S3 (B) when fermenting 20 g L^-1^ glucose, 80 g L^-1^ xylose, and 8 g L^-1^ acetate under micro-aerobic conditions. Data are presented as mean values and standard deviations of three independent biological replicates.

**Table 5.** Fermentation profiles of SR8A6S3-CDT_2_-GH43_2/7_, SR8A6S3-CDT_2_, and SR8A6S3 under anaerobic conditions.

|  | At 24 h |  |  |  | At 72 h |  |  |  |  |  |
| --- | --- | --- | --- | --- | --- | --- | --- | --- | --- | --- |
| | $r_{\text{xylose}}$ | $r_{\text{xylose}}^*$ | $P_{\text{xylitol}}$ | $P_{\text{Ethanol}}$ | $r_{\text{xylose}}$ | $r_{\text{xylose}}^*$ | $P_{\text{xylitol}}$ | $P_{\text{Ethanol}}$ | $Y_{\text{Ethanol}}$ | $Y_{\text{Xylitol}}$ |
| SR8A6S3-CDT<sub>2</sub>-GH43<sub>2/7</sub> | 0.54±0.03 | 0.15±0.01 | 0.00±0.00 | 0.77±0.05 | 0.57±0.01 | 0.10±0.00 | 0.01±0.00 | 0.45±0.00 | 0.47±0.01 | 0.018±0.000 |
| SR8A6S3-CDT<sub>2</sub> | 0.37±0.13 | 0.10±0.03 | 0.00±0.00 | 0.62±0.06 | 0.63±0.03 | 0.10±0.02 | 0.01±0.00 | 0.41±0.01 | 0.44±0.01 | 0.015±0.001 |
| SR8A6S3 | 0.94±0.06 | 0.19±0.01 | 0.00±0.00 | 0.74±0.00 | 0.83±0.02 | 0.14±0.00 | 0.03±0.00 | 0.47±0.00 | 0.42±0.00 | 0.032±0.0000 |

Parameters: $r_{\text{xylose}}$ , xylose consumption rate ( $\text{g L}^{-1} \text{h}^{-1}$ ); $r_{\text{xylose}}^*$ , specific xylose consumption rate ( $\text{g L}^{-1} \text{OD}^{-1} \text{h}^{-1}$ ); $P_{\text{xylitol}}$ , volumetric xylitol productivity ( $\text{g L}^{-1} \text{h}^{-1}$ ); $P_{\text{Ethanol}}$ , volumetric ethanol productivity ( $\text{g L}^{-1} \text{h}^{-1}$ ); $Y_{\text{Ethanol}}$ , ethanol yield ( $\text{g g}_{\text{consumed carbon source}}^{-1}$ ); $Y_{\text{Xylitol}}$ , xylitol yield ( $\text{g g}_{\text{consumed carbon source}}^{-1}$ ).

Concerning acetate metabolism, the control strain consumed 71% of the initial acetate in the medium, while SR8A6S3-CDT_2_ consumed only 43% in 72 h of cultivation (**Fig. 2.A**, **2.B,** and **S2.E**). For glucose metabolism, no difference was observed between the two strains (**Fig. 2**). However, despite the greater consumption of xylose and acetate by the control strain (83.08 ± 1.46 versus 70.58 ± 2.91 g L^-1^), SR8A6S3-CDT_2_ had a slightly higher ethanol yield (**Table 5**) and produced 66% less xylitol and 12% less glycerol as a by-product (**Fig. S2**). We observed that glycerol was primarily coming from glucose for both cultivations. Great amount of total glycerol was produced at 24 h of cultivation, 66% for SR8A6S3 and 60% for SR8A6S3-CDT_2_. Considering these results, it is possible to conclude that in the control strain cultivation, the carbon source was channelled towards metabolites whose pathways allowed the balance of redox cofactors, such as xylitol and glycerol. Thereby, *sor1* is responsible for a significant amount of xylitol production but *sor1*Δ primarily slows down xylose metabolism. Whereas *sor1*Δ enabled the engineered strain to drive more carbon toward the desired product (ethanol). Presumably, this is because the NADH / NAD^+^ balance has changed while the ethanol yield has increased marginally via the pyruvate decarboxylase (PDC) route as, relative to xylose, acetate metabolism is proportionally lower when the two strains are compared.

Some metabolites were measured to compare the fermentation profiles of SR8A6S3-CDT_2_ and SR8A6S3 (**Fig. S2**). Elimination of xylitol production through *sor1*Δ increases the availability of intracellular NADH, which enabled the recombinant cell to produce more ethanol per gram of consumed sugar (ethanol yield). Deletion of the *sor1* gene activity does not eliminate xylitol production since other genes encode enzymes capable of xylose reduction or xylitol oxidation, resulting in xylitol production. However, under strict anaerobic cultivation, the xylitol amount was reduced from 1.9 g L^-1^ to 0.69 g L^-1^ comparing parental and recombinant strains, respectively (**Fig. S2.C**). In principle, NAD^+^ should be available to drive the xylitol to xylulose reaction.

Under strict anaerobic conditions, ethanol is the most important primary metabolite produced in terms of re-oxidation of excess NADH and redox balancing, followed by the production of glycerol (Jain et al., 2011), which is important to support xylulose production from xylitol. When oxygen is available in the flask, redox balancing of NADH / NAD^+^ can also occur through the electron transport chain, which should result in less xylitol accumulation in the medium. We corroborated this hypothesis during batch cultivations under micro-aerobic conditions, where lower xylitol production was observed for both strains (**Fig. 3** and **S3.C**). Micro-aerobic batch fermentations were performed in complex YP media supplemented with 20 g L^-1^ glucose, 80 g L^-1^ xylose, and 8 g L^-1^ acetate with an initial OD_600_ of 1 (**Fig. 3** and **S3**).

Under micro-aerobic conditions the ethanol yields of SR8A6S3-CDT_2_ and SR8A6S3 were 0.39 g_Ethanol_ (g_consumed sugars_) and 0.37 g_Ethanol_ (g_consumed sugars_), respectively. As expected, xylitol yield was lower in SR8A6S3-CDT_2_ than in SR8A6S3, 0.004 gXylitol (g_consumed xylose_)^-1^ against 0.009 gXylitol (g_consumed xylose_)^-1^, respectively. Until 48 h of cultivation, SR8A6S3-CDT_2_ consumed 79% of the initial concentration of acetate, whereas SR8A6S3 consumed slightly lower amounts, 73%. After 48 h, both strains star oxidising the ethanol back to acetate (**Fig. 3**).

### 3.2. Cas9-based integration of a GH43-2_GH43-7 expression cassette into the *GRE3 locus*

GRE3 is an important xylose-reducing enzyme expressed by *S. cerevisiae* strains, the deletion of which decreases xylitol formation (Träff et al., 2001). Therefore, to further decrease carbon diverted to xylitol formation, a cassette for *GH43-2* and *GH43-7* high expression was integrated into the *gre3* locus of SR8A6S3-CDT_2_ using a CAS-9-based system (Stovicek et al., 2015), yielding the SR8A6S3-CDT_2_-GH43_2/7_ strain. The desired integration of the GH43_2/7_ sequence cassette into SR8A6S3-CDT_2_-GH43_2/7_ was confirmed by colony PCR performed on 7 colonies from the positive-control plate (**Fig. S4**). Once correct integration was confirmed, a positive transformant was then evaluated for growth in xylose and acetate, hydrolysed xylan, and hemicellulosic hydrolysate.

Thereby, to investigate the latest engineered strain, a YP-based medium was used to cultivate SR8A6S3-CDT_2_-GH43_2/7_ and measure xylose and acetate fermentation performance compared with SR8A6S3-CDT_2_ and their parental strain, SR8A6S3. Anaerobic batch cultivation was carried out for xylose and acetate consumption evaluation, and ethanol and xylitol production in high sugar content media (20 g L^-1^ glucose, 80 g L^-1^ xylose, and 8 g L^-1^ acetate) with an initial OD600 of 1 (**Fig. 2** and **S5**).

In the first 24 h of cultivation, SR8A6S3-CDT_2_-GH43_2/7_ had an increased xylose consumption profile, compared with the SR8A6S3-CDT_2_ strain. The latest engineered strain consumed 13.65 ± 0.53 g L^-1^ of xylose, which represents 18% of the initial xylose concentration, and the immediate parent consumed 10.73 ± 0.53 g L^-1^ (15% of the initial xylose concentration). During the same period, the acetate consumption profile was slightly higher for SR8A6S3-CDT_2_-GH342/7 than SR8A6S3-CDT_2_, 17% against 15% of the initial acetate concentration, respectively (**Fig. 2B** and **2C**). Following a similar line of analysis, the glycerol production profile, within the first 24 h, was higher for SR8A6S3-CDT_2_-GH342/7 than for SR8A6S3-CDT_2_ strain. The first produced 1.47 ± 0.04 g L^-1^ and the second one achieved 0.98 ± 0.13 g L^-1^ of glycerol. Conversely, within 24 and 72 h of cultivation, SR8A6S3-CDT_2_-GH43_2/7_ consumed lesser amounts of xylose and acetate than its immediate parent strain, 28.84 ± 0.66 g L^-1^ against 34.61 ± 3.45 g L^-1^ for xylose and 1.81 ± 0.14 g L^-1^ against 2.38 ± 0.14 g L^-1^ for acetate, respectively; as well as produced lesser amounts of glycerol, 0.34 ± 0.04 g L^-1^ against 0.66 ± 0.12 g L^-1^, respectively for SR8A6S3-CDT_2_-GH43_2/7_ and SR8A6S3-CDT_2_ (**Fig. 2B** and **2C**). The change in the profile of xylose, acetate, and glycerol for both strains in the first 24 h of cultivation and after this time, presumably is because of the change in the balance of NADH / NAD^+^. Deletion of *gre3* and increased production of glycerol (within 24 h of cultivation) might result in higher availability of the cofactors required for xylose metabolism (**Fig. S6**), which reflected better xylose consumption profile for SR8A6S3-CDT_2_-GH43_2/7_ in the first 24 h of cultivation. After the depletion of glucose, the glycerol production profile decreased for both strains (**Fig. 2B** and **2C**) but *gre3*Δ slows down xylose metabolism.

Instead, compared with SR8A6S3 (**Fig. 2A** and **2B**), the latest engineered strain had impaired xylose and acetate consumption profiles during all times of cultivation. Within the first 24 h of cultivation, SR8A6S3 consumed 22.53 ± 1.24 g L^-1^ of xylose, which represents 28% of the initial concentration, and 2.28 ± 0.10 g L^-1^ of acetate. The doubly engineered strain after 72h consumed only 56% and 40% of the initial concentration of xylose and acetate, respectively, although, intriguingly, rate of xylose consumption was marginally higher than the parent strains after 24h. However, SR8A6S3-CDT_2_-GH43_2/7_ had a slightly higher ethanol yield compared to both SR8A6S3-CDT_2_ and SR8A6S3 (**Table 5**). Therefore, although xylitol production after 72h was similar for SR8A6S3-CDT_2_-GH43_2/7_ and SR8A6S3-CDT_2_ deletion of both *sor1*Δ and *gre3*Δ,which should increase the availability of intracellular NADH and NADP^+^, enabled cells to produce more ethanol per gram of consumed sugar (ethanol yield) than the *sor1*Δ single deletion (**Fig. S6**). Moreover, the biomass production profile, which was analysed by measurement of OD_600_, of SR8A6S3-CDT_2_-GH43_2/7_ and SR8A6S3-CDT_2_ was similar until 24 h but was lower for the reference strain. The double-engineered strain and its immediate parent presented an increase in biomass content of 2.95 ± 0.01 and 2.81 ± 0.01, representing an increase of 376% and 384% of OD_600_ within the first 24 h of cultivation. In the meantime, SR8A6S3 achieved the growth of 584% of initial cell concentration, achieving at 24 h of cultivation an OD_600_ of 3.90 ± 0.03. The *sor1*Δ decreased the xylose consumption rate, while *gre3*Δ increased this rate at 24 h, but still lower than SR8A6S3 (**Table 5**).

Wenger et al. (2010) screened a large number of *S. cerevisiae* strains from wild, industrial, and laboratory backgrounds to determine the xylose-positive phenotype. Of 647 studied strains, some wine strains appeared to be able to grow modestly on xylose. By the application of high-throughput sequencing to bulk segregant analysis, they were able to identify a novel *XDH* gene homologous to *SOR1* (which was called *XDH1*) responsible for this phenotype. Next, the authors performed a comprehensive analysis of the involvement of the genes *GCY1, GRE3, YDL124W, YJR096W, YPR1, SOR1, SOR2, XDH1, XYL2*, and *XKS1* in the *XDH1* background strain (which has a xylose-positive phenotype) by single or combined deletion of the target genes. Single deletion of putative xylitol dehydrogenases (*SOR1, SOR2*, and *XYL2*) increased xylose utilization rate relative to the positive control; this phenotype was further enhanced when all three genes were deleted (*sor1*Δ *sor2*Δ *xyl2*Δ) (Wenger et al., 2010).

To assess the effect of endogenous xylitol-assimilating pathway genes on xylitol production profile by an engineered *S. cerevisiae* industrial strain CK17 overexpressing *Candida tropicalis XYL1* (encoding xylose reductase) in both batch and fed-batch fermentation with xylose and glucose as carbon sources, Yang et al. (2021) performed single deletion of the following genes: *XYL2* (yielding the strain *CK17Δxyl2), SOR1/SOR2* (yielding the strain CK17Δ*sor*), and *XKS1* (yielding the strain *CK17Δxks1*) (Yang et al., 2021). According to the authors, the mutant *sor*Δ had a reduced xylose consumption rate (12.4%) and xylitol production rate (4.7%), compared with its parental strain CK17, which is consistent with our findings for SR8A6S3-CDT_2_. The strain CK17Δ*xks1* had the highest xylose consumption rate (0.65 g L^-1^ h^-1^) and xylitol production rate (0.644 g L^-1^ h^-1^), while the control strain consumed xylose and xylitol at 0.598 g L^-1^ h^-1^ and 0.549 g L^-1^ h^-1^, respectively (Yang et al., 2021).

The *GRE3* gene was also deleted to improve xylose metabolism in *S. cerevisiae* CEN.PK2-1C expressing the xylose isomerase encoding gene *xylA* from *Thermus thermophilus*. The recombinant *gre3*Δ strains produced less xylitol than the parental strain (Träff et al., 2001). According to the authors, deletion of *GRE3* in *S. cerevisiae* decreased xylitol formation two- to threefold but not completely as xylitol may also be formed by the products of other genes, such as *XDH* (homologous to *SOR1* gene), through the reduction of xylulose or putative XR enzyme (Patiño et al., 2019; Richard et al., 1999; Wenger et al., 2010). Similarly, in the construction of a *S. cerevisiae* strain expressing the isomerase pathway (*xylA*) from the anaerobic fungus *Orpinomyces* sp. (GenBank No. MK335957), *gre3*Δ, *sor1*Δ, *XYL3*, and *TAL1* were added to reduce xylitol accumulation and increase the growth rate (Jeong et al., 2020).

On the other hand, overexpression of the endogenous genes *GRE3* and *XYL2*, coding for nonspecific aldose reductase and xylitol dehydrogenase, respectively, under endogenous promoters, enhanced the growth of *S. cerevisiae* on xylose in the presence of glucose in aerobic shake flask cultivation (Toivari et al., 2004). However, significantly more xylitol was formed by the CEN.PK2 strain overexpressing the *S. cerevisiae* enzymes in comparison to the strain that carries *XR* and *XDH* from *S. stipitis*. Also, transcriptional analysis of xylose and glucose grown cultures shows that the expression of *SOR1*, which encodes sorbitol dehydrogenase, was elevated in transformed cultures. Thereby, the presence of xylose resulted in higher *XDH* activity and induced the expression of the *SOR1* gene which also has *XDH* activity (Toivari et al., 2004).

*GRE3* and *SOR1* genes were considered for improving xylose fermentation based on these previous studies. In some of them, *sor1*Δ increased xylose utilization, and *gre3*Δ plus *sor1*Δ decreased xylitol accumulation. Similarly, we have observed that *gre3*Δ plus *sor1*Δ in *S. cerevisiae* SR8A6S3 decrease xylitol formation (**Table 5**). However, in contrast, *sor1*Δ alone did not increase the xylose consumption rate by SR8A6S3 (**Table 5**) as reported by (Wenger et al., 2010).

### 3.3. GH43 beta-xylosidases are intracellularly active

The activity of GH43-2 and GH43-7 in cell extracts of SR8A6S3 and SR8A6S3-CDT_2_-GH43_2/7_ was determined with pNPX as substrate (**Fig. 4**). No β-xylosidase activity was detected in the control strain, which is consistent with the absence of both genes *gh43-2* and *gh43-7* in its genome. On the other hand, the strain SR8A6S3-CDT_2_-GH43_2/7_ showed β-xylosidase activities of 27.74 U mL^-1^, or 114.53 U g_CDW_^-1^, or 0.160 U mg_protein_^-1^. The expression of both GH43-2 and GH43-7 is essential for converting XOS into xylose as the XR also acts as an XOS reductase, producing xylosyl-xylitol as a potential dead-end product, as first presented by Li et al., (2015). According to their work, despite the β-xylosidase GH43-7 having weak β-xylosidase activity, it rapidly cleaves xylosyl-xylitol into xylose and xylitol (Li et al., 2015).

**Fig. 4.**
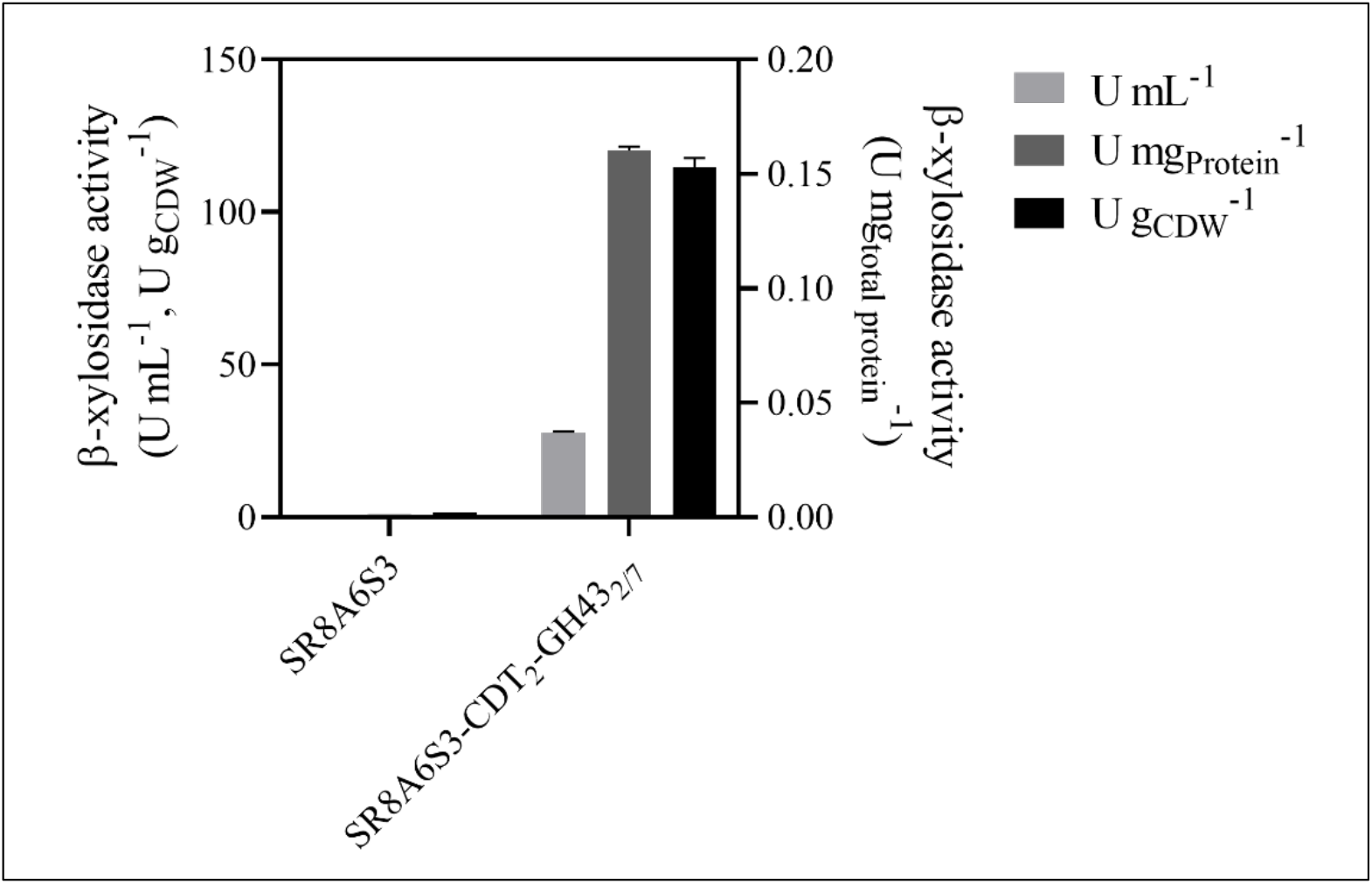
Intracellular β-xylosidase activity of SR8A6S3 and SR8A6S3-CDT_2_-GH43_2/7_ pellet extracts. The strains were cultured in YP-medium supplemented with 20 g L^-1^ glucose, 80 g L^-1^ xylose, and 8 g L^-1^ acetate) under microaerobic conditions until the late log phase. The intracellular GH43-2 and GH43-7 activities with pNPX as substrate were calculated relative to mg of protein and g of cell dry weight.

Within the context of XOS-to-ethanol production, other fungal xylanases have also been functionally expressed in *S. cerevisiae*, for example, β-xylosidase from *Aspergillus oryzae* NiaD300 and xylanase II from *Trichoderma reesei* QM9414, which had activities in *S. cerevisiae* MT8-1 of 234 U gCDW^-1^ and 16 U gCDW^-1^, respectively (Katahira et al., 2004); β-xylosidase from *T. reesei* QM9414 gave an activity of 6 nmol min^-1^ mg_Protein_^-1^ in *S. cerevisiae* M4-D4 (Fujii et al., 2011); Sakamoto and co-authors (2012) expressed an endoxylanase (*T. reesei*) and a β-xylosidase (*A. oryzae*) in *S. cerevisiae* MT8-1 and their activities were 41.2 U gCDW^-1^ and 16.8 U gCDW^-1^, respectively (Sakamoto et al., 2012); and most recently, endoxylanase from *T. reesei* QM6a was expressed in *S. cerevisiae* EBY100 giving activity of 1.197 U mg^-1^ (Tabañag et al., 2018).

### 3.4. Fermentation of hydrolysed xylan by the engineered SR8A6S3-CDT_2_-GH43_2/7_ strain

To evaluate XOS utilization, strain SR8A6S3-CDT_2_-GH43_2/7_ and the parental (control) strain SR8A6S3 were cultivated under micro-aerobic conditions at 30 °C, in a YP medium supplemented with hydrolysed xylan (YPXyl) and a mix of hydrolysed xylan plus acetate (YPAXyl) media. These media were designed to mimic a hemicellulosic hydrolysate but without the presence of inhibitory compounds, which can negatively influence yeast fermentations (Cola et al., 2020; Kłosowski and Mikulski, 2021). The engineered strain and its parental strain were cultivated in YPXy (**Fig. 5B** and **5D**), and in YPAXyl (**Fig. 5A** and **5C**) with an initial OD600 of 10 and, as expected, the engineered strain, SR8A6S3-CDT_2_-GH43_2/7_, produced higher titers of ethanol than the parental strain SR8A6S3 in all conditions tested.

**Fig. 5.**
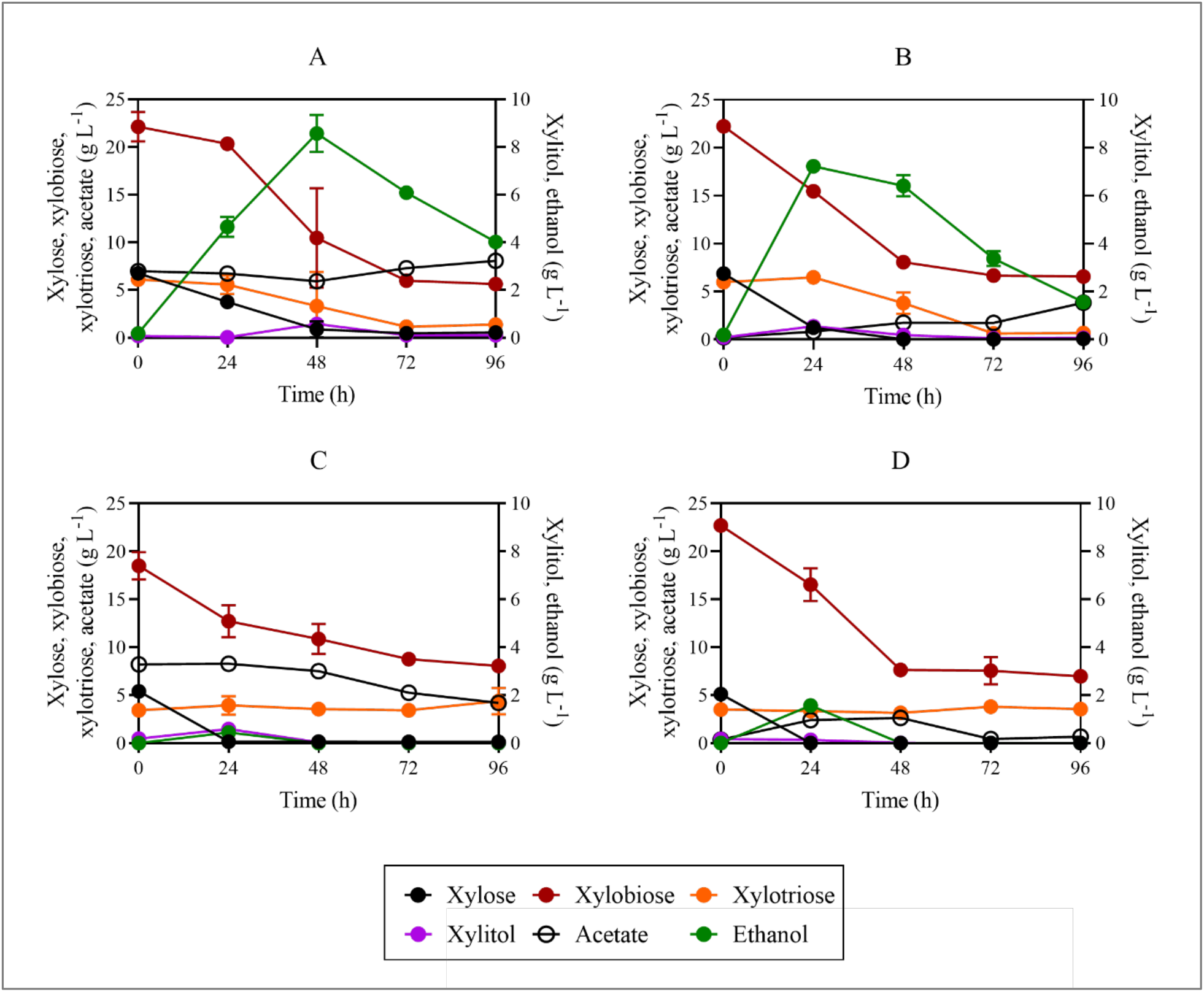
Fermentation profiles of SR8A6S3-CDT_2_-GH43_2/7_ (A and B) and SR8A6S3 (C and D) during batch cultivation in YPAXyl (YP medium containing hydrolysed xylan and acetate), A and C, and YPXyl (YP medium containing hydrolysed xylan), B and D. Cultivations were performed at 30 °C and 100 rpm with an initial OD600 of 10. Data are presented as the mean value and standard deviation of two independent biological replicates.

Xylobiose (X2) and xylotriose (X3) were the main carbon sources available in the medium. X2 concentrations decreased during the growth of both strains, SR8A6S3-CDT_2_-GH43_2/7_, and SR8A6S3 (**Fig. 5**), although SR8A6S3 did not express either heterologous xylanolytic enzymes or an XOS-transporter. One explanation could be that X2 entered the cell through a natural transport system in *S. cerevisiae* and was converted into the non-metabolizable compound xylosyl-xylitol by XR (xylose reductase), as observed previously by Li and colleagues (Li et al., 2015). It is important to note that *S. cerevisiae* can consume disaccharides such as maltose, sucrose, and trehalose, which are up taken through the action of membrane transporters (Lagunas, 1993). The uptake of sucrose (disaccharide composed of glucose and fructose) can occur via the proton-symport (*Mal11p*) (Marques et al., 2018). While trehalose (disaccharide composed of two glucose) can be taken up via Agt1p-mediated trehalose transport followed by intracellular hydrolysis catalysed by trehalase *Nth1*. Further, *AGT1/MAL11* gene is controlled by the *MAL* system. Maltose is transported to the cytosol by an energy-dependent process coupled to the electrochemical proton gradient (Lagunas, 1993).

Within the first 24 h of cultivation, SR8A6S3 depleted all xylose present in the medium (**Fig. 5C** and **5D**) while SR8A6S3-CDT_2_-GH43_2/7_ spent more time fermenting xylose completely (**Fig. 5A** and **5B**). At the same time, the doubly engineered strain consumed 6.77 ± 0.03 g L^-1^ of X2, which represents 30% of the initial X2 concentration and 1.79 ± 1.08 g L^-1^ of X2 (8% of the initial X2 concentration) respectively for the cultivations in YPXyl and YPAXyl. The presence of acetate changed the X2 consumption profile by SR8A6S3-CDT_2_-GH43_2/7_ (**Fig. 5A**). Intriguingly, the presence of X2 changed the acetate consumption profile by the parent strain, which consumed 4% of the initial acetate concentration until 24 h and 10% of the initial acetate concentration within first 48 h of cultivation (**Fig. 5C**). The slight acetate reduction within 24 and 48 h of cultivation might be affected by the oxidation of ethanol (Xu et al., 2022), which peak was at 24 h (**Fig. 5C**). Concerning the X3 consumption profile, the parent strain SR8A6S3 barely metabolized X3 in either medium (**Fig. 5C** and **5D**). Conversely, the engineered strain started to metabolize X3 after 24 h. The higher initial concentration of X2 than X3 probably interfered in X3 transportation. Instead, in 24 – 48 h, SR8A6S3-CDT_2_-GH43_2/7_ consumed 2.67 ± 0.85 g L^-1^ and 1.73 ± 1.32 g L^-1^ of X3 from YPXyl (**Fig. 5B**) and YPAXyl (**Fig. 5A**) cultivations, respectively. After 72 h of cultivation, no substantial decrease in X3 amount was observed for XOS-consuming strain cultivations.

The ethanol yield from SR8A6S3-CDT_2_-GH43_2/7_ was much higher than the control in both media (**Table 6**). Furthermore, although in 96 h of cultivation both strains consumed approximately the same amount of X2 in YPXyl (**Fig. 5B** and **5D**), only strain SR8A6S3-CDT_2_-GH43_2/7_ appeared to ferment it to ethanol. Deletion of *gre3* and *sor1* delayed xylitol production by SR8A6S3-CDT_2_-GH43_2/7_ strain. Interestingly, similar amounts of xylitol were observed for both control and engineered strains when cultured in YPAXyl, 0.59 ± 0.00 g L^-1^ and 0.57 ± 0.10 g L^-1^, respectively. However, at 24 h of cultivation for SR8A6S3 and 48 h for SR8A6S3-CDT_2_-GH43_2/7_.

**Table 6.**
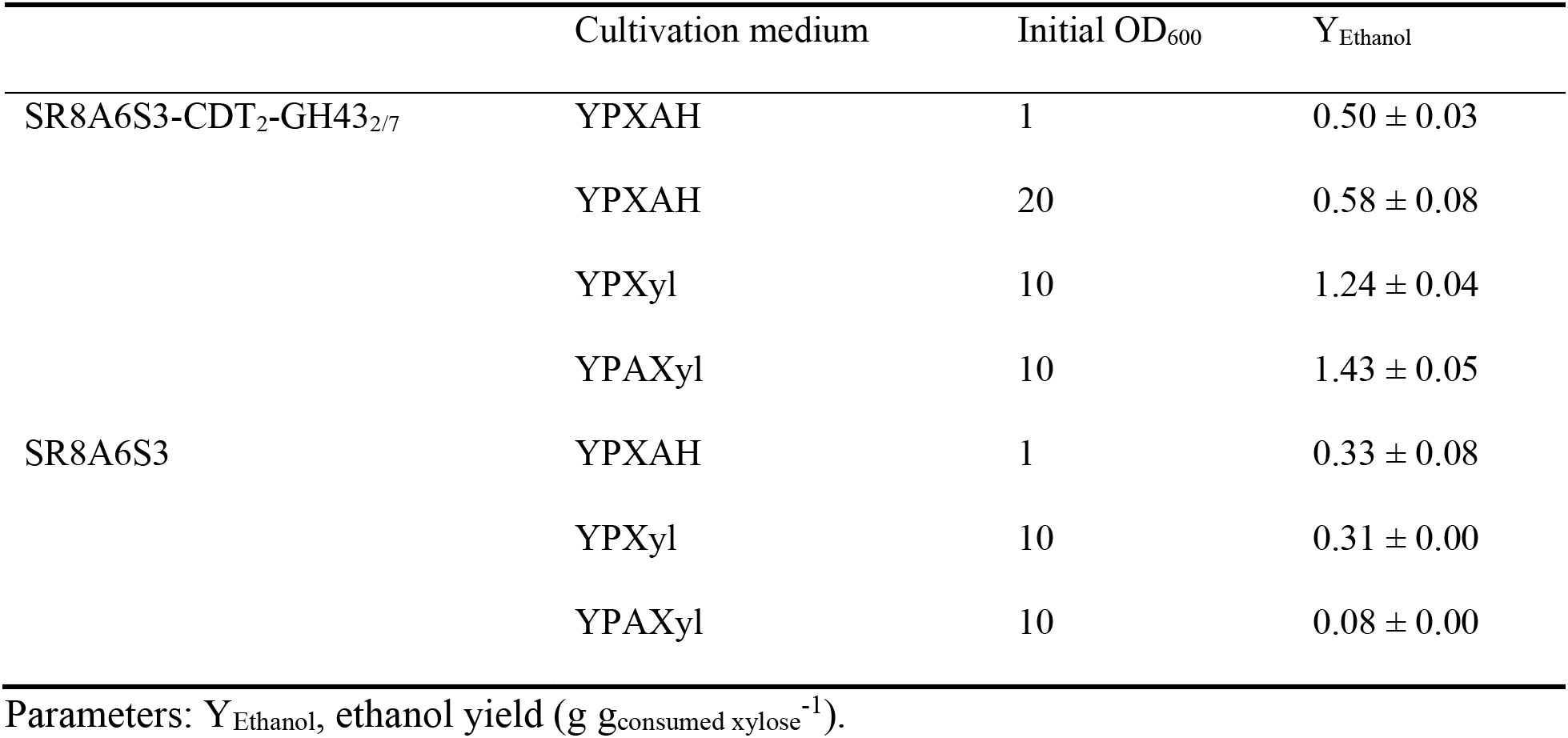
Ethanol yield of SR8A6S3-CDT_2_ and SR8A6S3 under micro-aerobic cultivation at 30 °C, in YP medium, supplemented with a mix of hemicellulosic hydrolysed plus xylose and acetate (YPXAH), hydrolysed xylan (YPXyl) or a mix of hydrolysed xylan plus acetate (YPAXyl) with varied initial OD_600_.

|  | Cultivation medium | Initial OD<sub>600</sub> | Y<sub>Ethanol</sub> |
| --- | --- | --- | --- |
| SR8A6S3-CDT<sub>2</sub>-GH43<sub>2/7</sub> | YPXAH | 1 | 0.50 ± 0.03 |
|  | YPXAH | 20 | 0.58 ± 0.08 |
|  | YPXyl | 10 | 1.24 ± 0.04 |
|  | YPAXyl | 10 | 1.43 ± 0.05 |
| SR8A6S3 | YPXAH | 1 | 0.33 ± 0.08 |
|  | YPXyl | 10 | 0.31 ± 0.00 |
|  | YPAXyl | 10 | 0.08 ± 0.00 |

Parameters: Y<sub>Ethanol</sub>, ethanol yield (g g<sub>consumed xylose</sub><sup>-1</sup>).

### 3.5. Fermentation of hemicellulosic hydrolysate by the engineered SR8A6S3-CDT_2_-GH43_2/7_ strain

Following successful cultivation in a simulated hemicellulose hydrolysate, strain SR8A6S3-CDT_2_-GH43_2/7_ was cultivated under micro-aerobic conditions in a YP medium supplemented with an authentic XOS-rich hemicellulosic hydrolysate (Brenelli et al., 2020) (**Fig. 6A** and **6B**), which mimics the context of a lignocellulosic biorefinery, which makes full use of hemicellulose. The breakdown of hemicellulose, which is acetylated (Kłosowski and Mikulski, 2021) releases highly toxic acetate, reducing the fermentative performance of *S. cerevisiae* (Bellissimi et al., 2009; Li et al., 2015). SR8A6S3 was previously engineered through an optimized expression of *AADH* and *ACS* in the acetate reduction pathway, enabling acetate conversion into ethanol by the optimized strain (Zhang et al., 2016). We, therefore, tested whether the acetate reduction pathway could operate simultaneously with XOS fermentation, as a means to augment ethanol yield from the lignocellulosic hydrolysate.

**Fig. 6.**
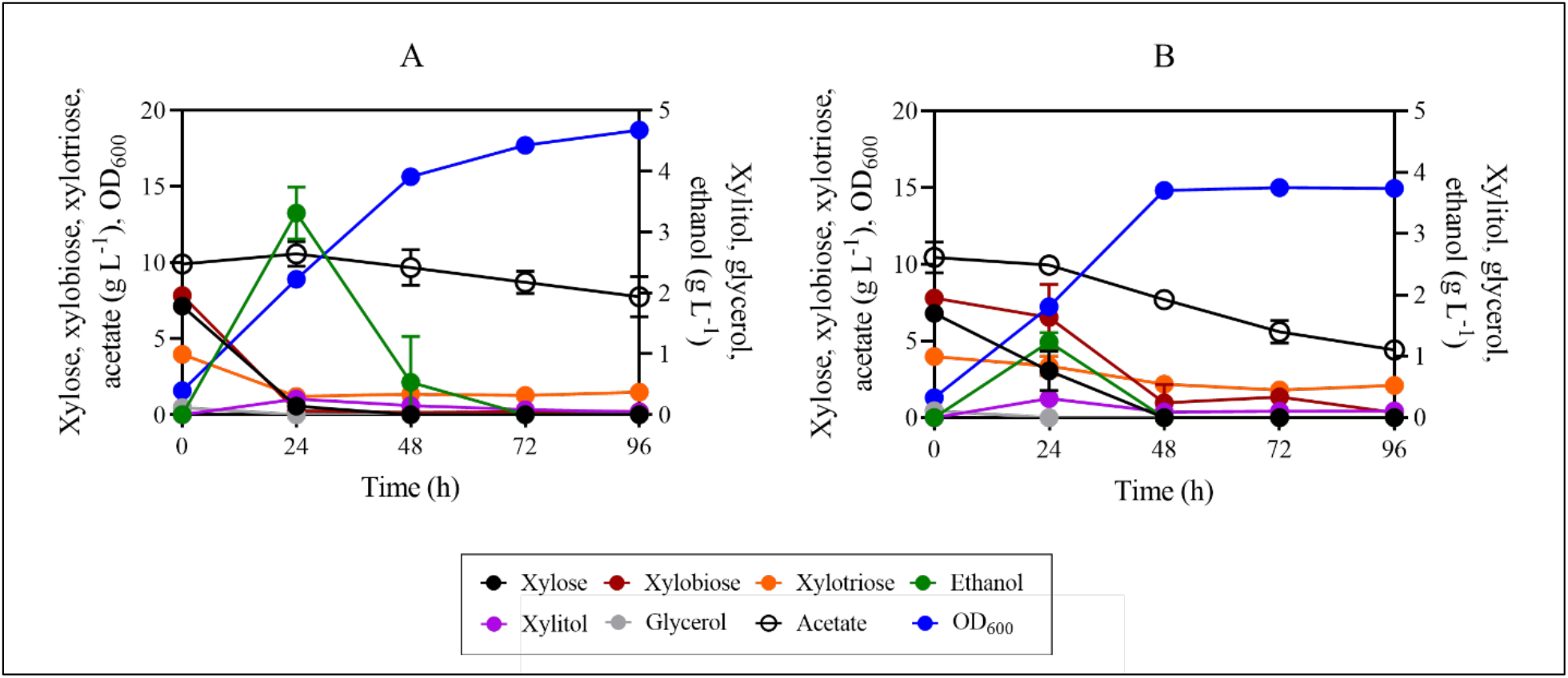
Fermentation profiles of SR8A6S3-CDT_2_-GH43_2/7_ (A) and SR8A6S3 (B) during batch cultivation in YPXAH (YP medium containing xylose, acetate, and hydrolysed hemicellulose). Cultivations were performed at 30 °C and 100 rpm with an initial OD600 of 1. Data are presented as mean values and standard deviations of two independent biological replicates.

Under this condition, we observed that xylose, X2, and X3 presented similar consumption profiles in the XOS-consuming strain cultivation. These carbon sources were primarily consumed before 24 h of cultivation. The latest engineered strain consumed 6.57 ± 0.28 g L^-1^ of xylose, which represents 92% of the initial xylose concentration, and 7.57 ± 0.08 g L^-1^ of X2, which represents 97% of the initial X2 concentration, and 2.76 ± 0.14 g L^-1^ of X3 (69% of the initial X3 concentration). Conversely, during the same period, the parent strain consumed only 3.73 ± 0.94 g L^-1^ of xylose (55% of the initial xylose concentration), and 16% and 15.5% of the initial X2 and X3 concentrations, respectively. It is worth pointing out that, as abovementioned, SR8A6S3 did not express either heterologous xylanolytic enzymes or an XOS-transporter, the uptake of XOS probably occurs through the action of membrane transporters that carry out disaccharides transport.

Although we observed a decrease in X2 and X3 amounts in the cultivations with the control strain, only in cultivations with the XOS-consuming strain (SR8A6S3-CDT_2_-GH43_2/7_) ethanol accumulation was consistent with X2 fermentation, i.e., conversion of X2 into ethanol (**Fig. 6A**). SR8A6S3-CDT_2_-GH342/7 and SR8A6S3-CDT_2_ achieved the highest ethanol concentration at 24 h of cultivation, 3.78 ± 0.53 g L^-1^ and 1.23 ± 0.10 g L^-1^, respectively. In other terms, the newest engineered strain achieved an ethanol yield of 0.50 ± 0.03 g g_consumed xylose_^-1^ and for the control strain ethanol yield was 0.33 ± 0.08 g g_consumed xylose_^-1^ (**Table 6**). Interestingly, the ethanol peach did not follow xylose exhaustion in the control cultivation (**Fig. 6B**), as happened to the engineered strain cultivation (**Fig. 6A**).

Acetate consumption was not observed in both SR8A6S3 and SR8A6S3-CDT_2_-GH342/7 cultivations within the first 24 h of cultivation. These results might indicate that transportation of X2 and X3 might result in changes of the balance of NADH / NAD^+^ and ATP which impaired acetate consumption profile. In a previous study, Zhang et al. (2016) highlighted that three major factors might limit the metabolic fluxes of the acetate reduction pathway, which include the intracellular ATP levels, NADH levels, and the activities of key enzymes (*ACS* and *AADH*), being the last the major limiting factor among them. In this study, the expression of key enzymes was not modified through genetic interventions. In 24 – 96 h of cultivation, acetate was reduced by 25% and 53% for the XOS-consuming and control strains, respectively. The acetate profile seems to be a combination of acetate consumption and acetate production, resulting from ethanol oxidation (Xu et al., 2022). The lesser change in acetate consumption profile for SR8A6S3-CDT_2_-GH342/7 than SR8A6S3 might have resulted from the higher amount of ethanol produced by this strain, which could be converted into acetate after exhaustion of the sugars (Xu et al., 2022). Hence, it appears to use less acetate. Regarding xylitol production, the SR8A6S3-CDT_2_-GH43_2/7_ strain produced a lower xylitol yield, 0.041 ± 0.01 g g_consumed xylose_^-1^, than the control cultivation, in which the yield was 0.083 ± 0.01 g g_consumed xylose_ ^-1^.

To evaluate the improvement obtained by the introduction of the XOS-consumption pathway in the SR8A6S3 strain, ethanol yield based on grams of consumed xylose was calculated for each condition (**Table 6**). The ethanol yield of the SR8A6S3-CDT_2_-GH43_2/7_ strain increased substantially as compared to the SR8A6S3 strain. This substantial yield increase is very likely due to the conversion of XOS to ethanol.

Lignocellulose-derived ethanol provides environmental and economic benefits besides being a promising industry in the expected transition from fossil fuels to renewable energy (Kłosowski and Mikulski, 2021). Hemicellulosic-derived sugar comprises 15-35% of lignocellulosic biomass, representing a large source of renewable material that is available at a low cost (Dahlman et al., 2003; Gírio et al., 2010; Kłosowski and Mikulski, 2021). Engineered strains able to consume XOS derived from hemicellulose via intracellular hydrolysis represent a potential benefit for bioethanol production since these strains would have a competitive advantage concerning other microorganisms, such as contaminating bacteria and wild *Saccharomyces* and non-*Saccharomyces* species that are expected to be unable to utilize XOS as a carbon source (Amorim et al., 2011; Procópio et al., 2022).

## 4. CONCLUSIONS

Xylose metabolism to ethanol in *S cerevisiae* SR8A6S3 is metabolically inefficient due to the production of xylitol. In this study we have integrated genes necessary to create a XOS-consumption pathway into two xylitol-production-related genes, *SOR1* and *GRE3*. The resulting strains, SR8A6S3-CDT_2_ and SR8A6S3-CDT_2_-GH43_2/7_, which are *sor1*Δ and *sor1*Δ, *gre3*Δ, respectively, showed a reduction in xylitol production and improvement in ethanol yield when compared with their parental strain SR8A6S3 in YPDXA cultivations under both micro-aerobic and anaerobic conditions. However, this coincided with a reduced rate of xylose metabolism, implying that there is scope for improvement in overall flux from xylose to ethanol. SR8A6S3-CDT_2_-GH43_2/7_ was able to ferment X2 and X3 efficiently for ethanol production and achieved the highest apparent ethanol yield (based only on the content of monomeric xylose) of 1.43 ± 0.05 g g_consumed xylose_^-1^ (64% higher than theoretical ethanol yield) in YP supplemented with hydrolysed xylan and acetate. When grown on a medium containing hemicellulose hydrolysate with low monomeric xylose content, fermentation of X2 and X3 was poor, but this was dramatically improved by the addition of monomeric xylose. This, and other evidence which shows that X2 and X3 metabolism slows down once the monomeric carbohydrates have been depleted, suggests that the latter are required to provide the energy demands of the former (e.g. for enzyme biosynthesis). While there is clearly room for further improvement, this demonstrates that a XOS fraction generated by simple hydrothermal/steam explosion pre-treatment of lignocellulosic agricultural residues, without any subsequent enzymatic hydrolysis, is a potential resource for renewable biofuel production using a XOS-utilising yeast.

## 5 CREDIT AUTHORSHIP CONTRIBUTION STATEMENT

**Dielle P. Procópio:** Methodology, Investigation, Formal analysis, Data curation, Validation, Writing – original draft, preparation. **Jae W. Lee:** Methodology. **Jonghyeok Shin:** Methodology. **Robson Tramontina:** Methodology. **Fabio Squina:** Formal analysis **André Damasio:** Formal analysis, Resources, Writing – review & editing. **Lívia P. Brenelli:** Methodology and Formal analysis. **Sarita C. Rabelo:** Formal analysis. **Rosana Goldbeck:** Methodology and Formal analysis. **Telma T. Franco:** Resources and Formal analysis. **David Leak**: Formal analysis, Writing – review & editing. **Yong-Su Jin:** Supervision, Formal analysis, Project administration. **Thiago O. Basso:** Supervision, Formal analysis, Resources, Project administration, Writing – review & editing.

## 6. ACKNOWLEDGMENT

We would like to thank The Brazilian Biorenewables National Laboratory (LNBR/CNPEM/MCTIC) for infrastructure.

## 7. FUNDING

This work was supported by the São Paulo Research Foundation (FAPESP) [grants numbers #2015/50590-4, #2015/50612-8, #2017/15477-8, #2018/17172-2, #2018/01759-4, #2019/18075-3, and #2021/04254-3].

## Notes

### Competing Interest Statement

The authors have declared no competing interest.

