## Supplementary material for "Metabolic engineering of *Saccharomyces cerevisiae* for second-generation ethanol production from xylo-oligosaccharides and acetate"

**Table S1**. Primers used in this study.

| Name | Sequence (5'- 3') |
| --- | --- |
| DPO_059 | ATGCCCCTCGTCAAGAACCCCATCCTCCCCGGCTTCAATC |
| DPO_063 | TTACTTCCCAGCCGGCTGCTTTTCCCCACAAATCTTCCCCTCTTCA |
| DPO_064 | CCGGGCTGCAGGAATTCGAT |
| DPO_065 | GGGATCCACTAGTTCTAGAA |
| DPO_062 | CCAGAACTTAGTTTCGACGGATTCTAGAACTAGTGGATCCCATGCCCCTCGTCAAGAACC |
| DPO_074 | GACGGTATCGATAAGCTTGATATCGAATTCCTGCAGCCCGGTTACTTCCCAGCCGGCTGC |
| DPO_057 | GTAATATAAATCGTAAAGGAAAATTGGAAATTTTTTAAAGGTAATACGACTCACTATAGG |
| DPO_058 | TTGTTCATATCGTCGTTGAGTATGGATTTTACTGGCTGGAAATTAACCCTCACTAAAGGG |
| DPO_081 | AATCAACAAGAAAAAATACTAAAAAAAAAAAATTGAAAAATGTAAAACGACGGCCAGT |
| DPO_082 | TATATATGGACATGAACCAGTGCCGAAAAGTATTCACTTTACAGGAAACAGCTATGAC |
| DPO_089 | TGTGTCGAACCCTTATCAGTGTTTTAGAGCTAGAAATAGCAAG |
| DPO_090 | ACTGATAAGGGTTCGACACAGATCATTTATCTTTCACTGCGGA |
| DPO_083 | CCGGTCTCGTATCTCCTTT |
| DPO_084 | CTATCAACTGGAAGTAATGCG |
| DPO_069 | GGGGGCCTATCAAGTAAATTACTCCTGGT |
| DPO_070 | GTTCAGATTCACTTCTTGATATTTCC |
| DPO_087 | TCCTCAATCATTCATTGAGAGTTTTAGAGCTAGAAATAGCAAG |
| DPO_088 | TCTCAATGAATGATTGAGGAGATCATTTATCTTTCACTGCGGA |

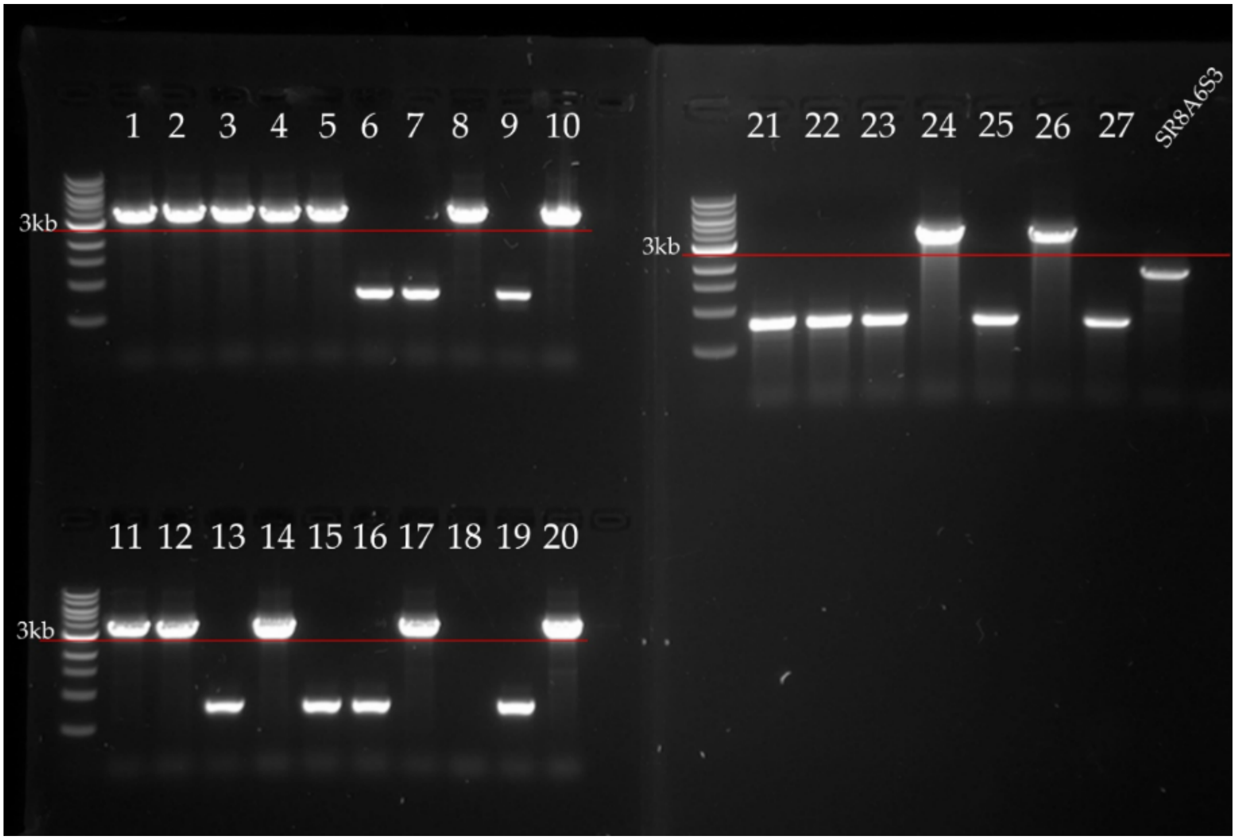


**Fig. S1.** Colony PCR products were analyzed using an agarose gel of 0.8% from positive control plate colonies. *sor1*Δ::$_{P}$*PGK1-CDT-2-*$_{T}$*CYC1* mutant cells present a band of 3.4 kb (1, 2, 3, 4, 5, 8, 10, 11, 12, 14, 17, 20, 24, 26), a deletion of the sor1 gene was observed in 6, 7, 9, 13, 15, 16, 19, 21, 22, 23, 25, 27 which band present 0.8 kb. The last band, SR8A6S3, represents the positive PCR control, of which the band has 1.8 kb.

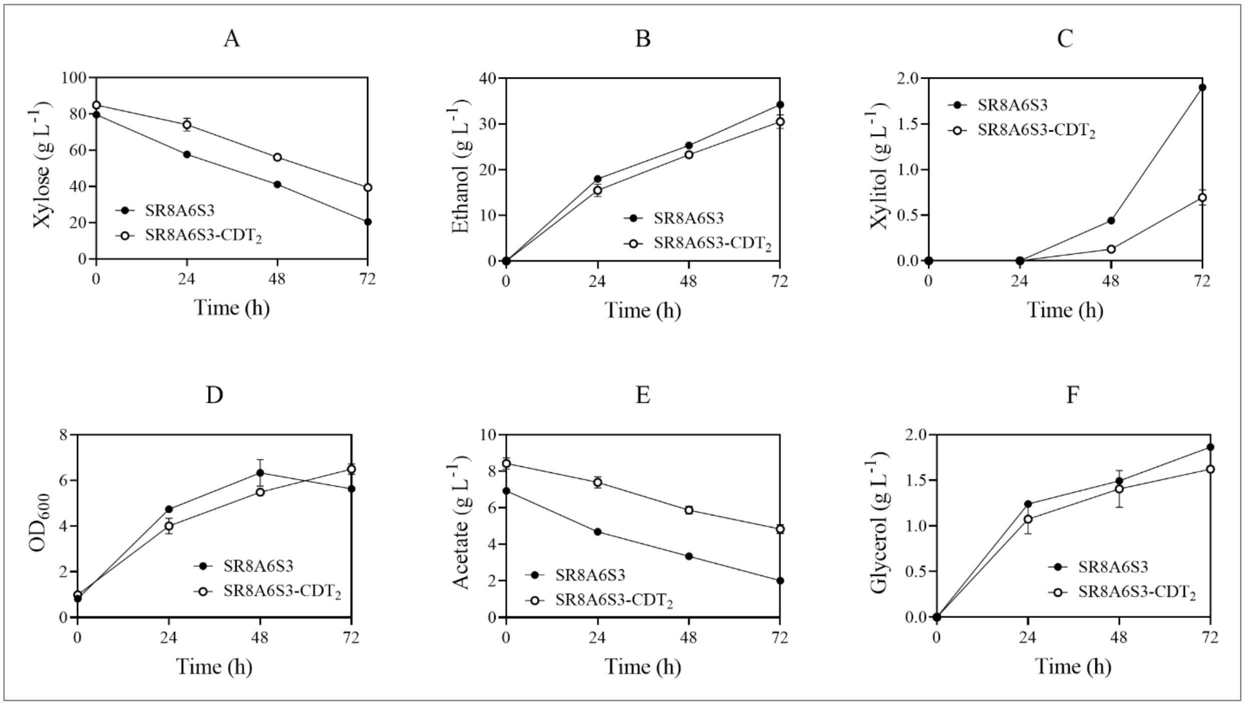


**Fig. S2**. Comparison of xylose (A) and acetate (E) consumption, ethanol (B), xylitol (C), $OD_{600}$ (D), and glycerol (F) production of two xylose-acetate-assimilating strains, SR8A6S3-$CDT_2$ and SR8A6S3, in YP media containing 20 g $L^{-1}$ glucose, 80 g $L^{-1}$ xylose, and 8 g $L^{-1}$ acetate under anaerobic condition. An initial $OD_{600}$ was adjusted to 1. The figure illustrates the means of triplicate experiments for each strain.

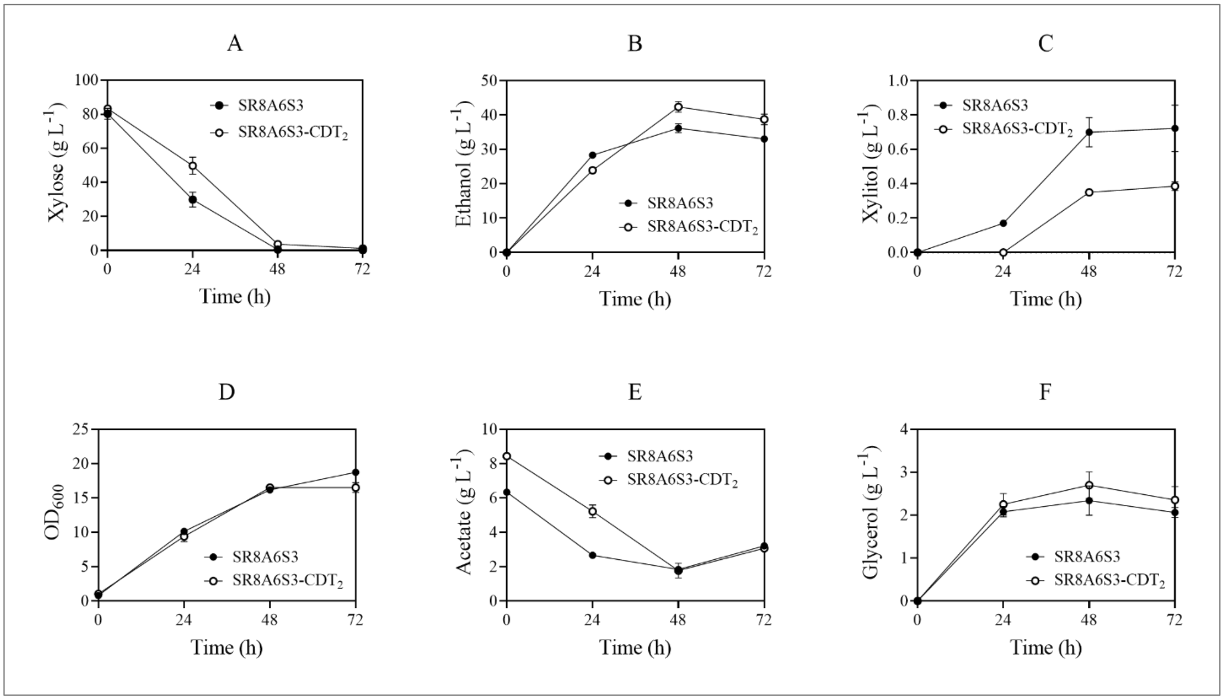


**Fig. S3**. Comparison of xylose (A) and acetate (E) consumption, ethanol (B), xylitol (C), $OD_{600}$ (D), and glycerol (F) production of two xylose-acetate-assimilating strains, SR8A6S3-$CDT_2$ and SR8A6S3, in YP media containing 20 g $L^{-1}$ glucose, 80 g $L^{-1}$ xylose, and 8 g $L^{-1}$ acetate under micro-aerobic condition. An initial $OD_{600}$ was adjusted to 1. The figure illustrates the means of triplicate experiments of each strain.

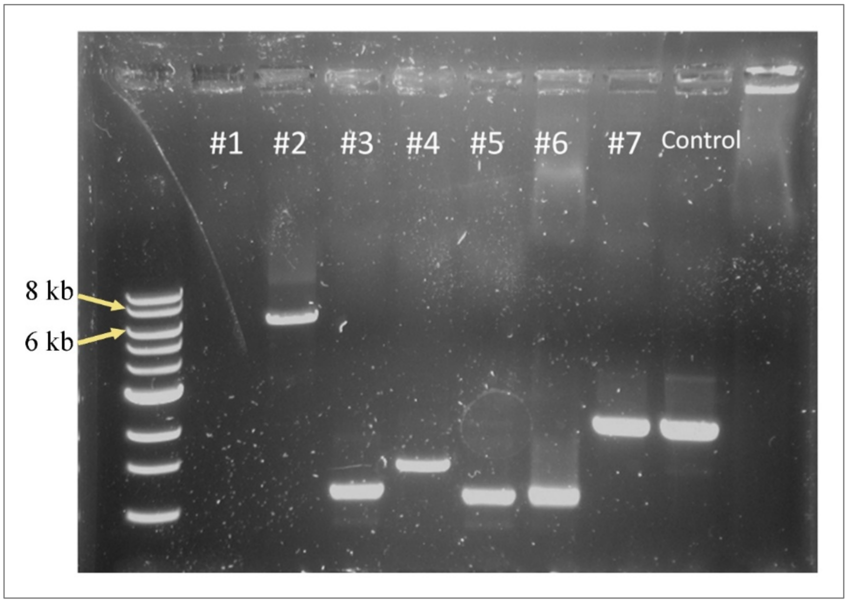


**Fig. S4**. Colony PCR products analyzed using an E-Gel 0.8% agarose from positive control plate colonies. *gre3*Δ:: $_{P}$TDH3-GH43-7-$_{T}$CYC- $_{P}$CCW12-GH43-2-$_{T}$CYC1 mutant cells present a band of 7 kb (colony 2), The last band represents the positive PCR control, of which the band has 1.5 kb.

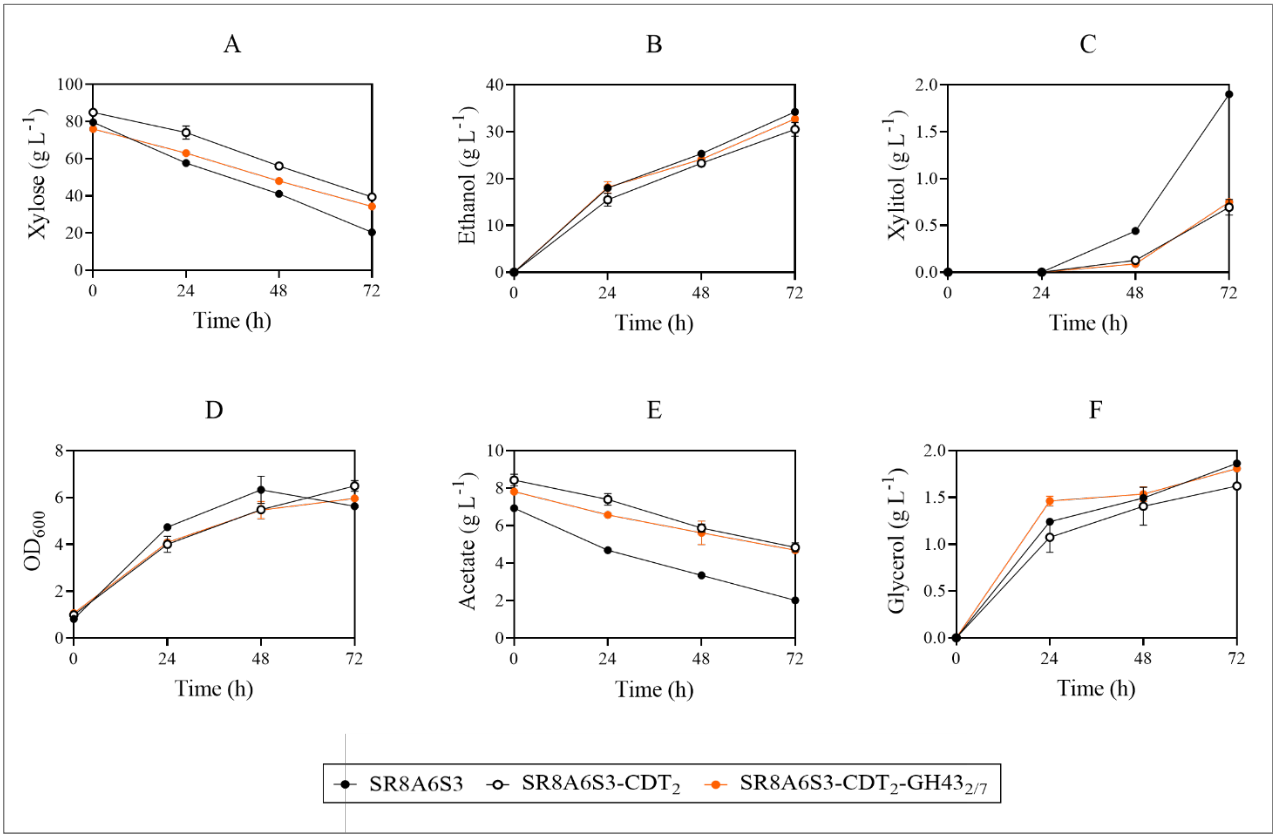


**Fig. S5**. Comparison of xylose (A) and acetate (E) consumption, ethanol (B), xylitol (C), $OD_{600}$ (D), and glycerol (F) production of three xylose-acetate-assimilating strains, SR8A6S3, SR8A6S3-$CDT_2$, and SR8A6S3-$CDT_2$-$GH43_{2/7}$ cultivated in YP media containing 20 g $L^{-1}$ glucose, 80 g $L^{-1}$ xylose, and 8 g $L^{-1}$ acetate under anaerobic condition. An initial $OD_{600}$ was adjusted to 1. The figure illustrates the means of triplicate experiments of each strain.

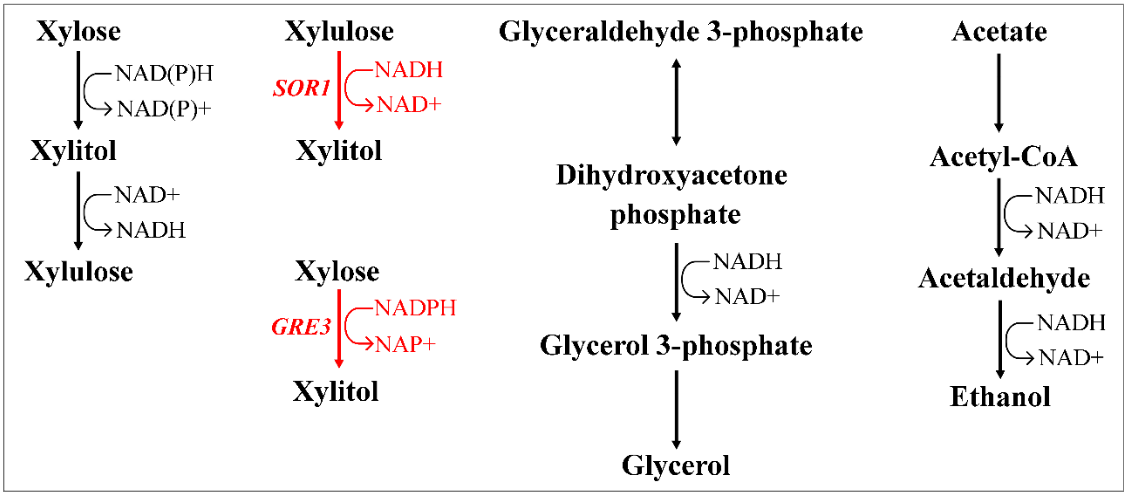


**Fig. S6**. Target metabolites were analyzed for comparison of fermentation profiles of the SR8A6S3-CDT$_2$-GH43$_{2/7}$ and SR8A6S3. The red arrow represents deleted gene/route.
